# Biological condensates form percolated networks with molecular motion properties distinctly different from dilute solutions

**DOI:** 10.1101/2022.07.20.500769

**Authors:** Zeyu Shen, Bowen Jia, Yang Xu, Jonas Wessén, Tanmoy Pal, Hue Sun Chan, Shengwang Du, Mingjie Zhang

## Abstract

Formation of membraneless organelles or biological condensates via phase separation hugely expands cellular organelle repertoire. Biological condensates are dense and viscoelastic soft matters instead of canonical dilute solutions. Unlike discoveries of numerous different biological condensates to date, mechanistic understanding of biological condensates remains scarce. In this study, we developed an adaptive single molecule imaging method that allows simultaneous tracking of individual molecules and their motion trajectories in both condensed and dilute phases of various biological condensates. The method enables quantitative measurements of phase boundary, motion behavior and speed of molecules in both condensed and dilute phases as well as the scale and speed of molecular exchanges between the two phases. Surprisingly, molecules in the condensed phase do not undergo uniform Brownian motion, but instead constantly switch between a confined state and a random motion state. The confinement is consistent with formation of large molecular networks (i.e., percolation) specifically in the condensed phase. Thus, molecules in biological condensates behave distinctly different from those in dilute solutions. This finding is of fundamental importance for understanding molecular mechanisms and cellular functions of biological condensates in general.

## Introduction

Phase separation-mediated formation of condensed macro-molecular assemblies is being recognized as a general mechanism for cells to form a distinct class of cellular organelles with diverse functions (Banani et al., 2017; Chen et al., 2020; Lyon et al., 2021; Shin et al., 2017; Wu et al., 2020). Compared to the classical cellular organelles that are demarcated by lipid membranes, organelles formed via phase separation either do not associate with or are not enclosed by lipid membranes and such organelles are referred to as biological condensates or membraneless organelles in the literature (we use biological condensates throughout this paper). Formation of biological condensates greatly expands the means of how a living cell compartmentalizes its molecular constituents for specific and diverse functions. Since biological condensates are not enclosed by membranes, molecules within a biological condensates are in dynamic exchange with the counterparts in dilute solution without energy input, thus establishing a foundation for numerous unique properties of biological condensates (e.g. how sharp concentration gradient between the condensed and dilute phases is maintained; how molecules are selected to be included or excluded in the condensates; means and rates to regulate condensate formation/dispersion; etc.) with respect to the membrane-enclosed organelles. The concept of biological condensate formation and function has gained extensive interests in recent years, but the field is still in its infancy and sometimes under debate (McSwiggen et al., 2019; Mittag and Pappu, 2022; Musacchio, 2022).

Molecules within biological condensates can be massively concentrated. For example, proteins can be concentrated by more than 10,000 folds upon chromatin condensate formation (Gibson et al., 2019). In cell peripherals such as synapses in neurons, phase separation can concentrate numerous proteins into postsynaptic densities by >1,000-fold (Zeng et al., 2018). A fundamental task in biological phase separation research is to understand how molecules in the condensed phase behave and function. The existing biochemistry and biophysics theories that have been guiding our understandings of molecular behaviours and their interactions in living cells in the past are mainly developed for molecules in dilute solutions. A biological condensate formed via phase separation is more of a condensed soft matter system, thus theories dealing with dilute solutions are not expected to be generally adequate for condensed molecular systems. Due to extreme complexities of molecular constituents (i.e., proteins, nucleic acids and lipids), molecular compositions (i.e., each functional biological condensate often contains hundreds or more different types of molecules), and broad range of interaction modes (e.g., very large dynamic ranges of binding affinities and molecular valency, different levels of cooperativities, etc.) of biological condensates in cells, currently available theories in soft matter physics and polymer chemistry, though very useful, are likely not sufficient to be directly adapted to characterize biological condensates. Experimental methods currently used to study molecules in biological condensates are largely qualitative and descriptive.

Here we develop an adaptive super-resolution imaging-based method that can simultaneously and robustly monitor and quantify motion properties of individual molecules in dilute and condensed phases of biological condensates formed in solution or on lipid membranes. In addition to directly visualizing motion trajectories within and between phases, this method affords direct measurements of diffusion parameters of each molecule in the dilute and condensed phase. Unexpectedly, we observed that molecules in the condensed phase spend a very large fraction of time in transient motion-frozen state. Such temporary motion freeze exists in various biological condensates and is governed by specific and multivalent interaction-mediated large molecular network formation in condensed phases. The motion property changes due to formation biological condensates can fundamentally alter action mechanisms and cellular functions of biomolecules.

## Results

### Localization-based super-resolution imaging of phase separation

In an earlier study, we introduced localization-based single molecule tracking experiment to study motion properties of proteins in the condensed phase of in vitro reconstituted active zone condensates formed on two-dimensional supported lipid bilayer (SLB) (Wu et al., 2019). Here, we further developed the method into an assay that can simultaneously track molecules in both condensed and dilute phases. We used the in vitro reconstituted postsynaptic density (PSD) condensates formed on SLB (Zeng et al., 2018) to demonstrate this method. Four major PSD proteins (PSD95, SHANK3, GKAP, Homer) and Trx-tagged GCN4-His_8_-NR2B-CT tetramer (termed as NR2B in this article) were included in our study (Figure 1A). These five proteins, via specific and multivalent interactions, form a large molecular network capable of phase separation at physiological concentrations (Zeng et al., 2018).

**Figure 1:**
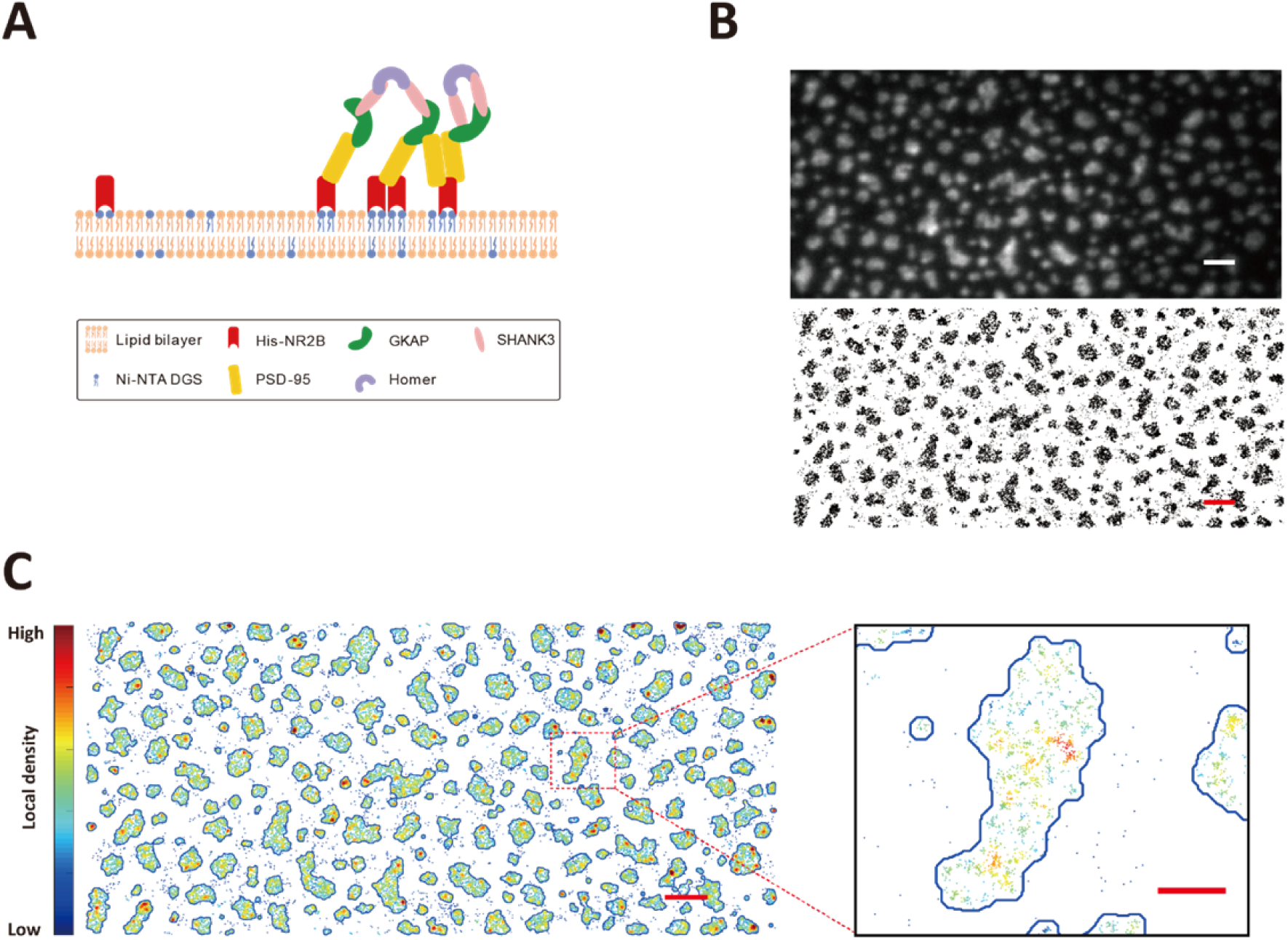
Single molecule imaging of phase separation on supported lipid bilayers. (A) Schematic diagram showing phase separation of PSD protein assembly on SLBs (Zeng et al., 2018). (B) Upper panel: a TIRF image of Alexa647 labelled His_8_-NR2B tetramer clustered within the PSD condensate on SLB. Lower panel: Stacking of 4000 frames of dSTORM images of Alexa 647 labelled His_8_-NR2B within the same PSD condensates as shown in the TIRF image above. Black dots represent localizations recognized during the imaging. Scale bar: 2 μm. (C) Phase boundary of the PSD condensates determined by localization densities. The boundaries are shown in blue lines. Localizations are color-coded according to their local densities from low (blue) to high (red). A zoom in view of a typical condensed patch on SLB, showing heterogeneous distributions and nano-cluster-like structures of molecules within the condensed phase. Scale bar of the original image: 2 μm, scale bar for the zoom in view: 500 nm.

Since the densities of proteins are hugely different between condensed and dilute phases, sparse labelling would lead to lack of information for molecules in the dilute phase and dense labelling would cause extensive overlapping of single molecule signals in the condensed phase during conventional fluorescence imaging experiments. To overcome this dilemma, we utilized dSTORM imaging (van de Linde et al., 2011) to obtain a large number of stochastically emitted single molecule tracks in both condensed and dilute phases by labelling proteins with photo-switchable dyes (in this case by labelling NR2B with 1% Alexa647). TIRF illumination mode was used to detect protein signals on SLB so that signals from molecules not tethered to the membrane were minimized.

The PSD mixtures formed noncircular condensed phase on SLB with around or less than 1 μm in size, but conventional TIRF images could only provide fuzzy phase boundaries at this scale (Figure 1B, *top*; also see (Zeng et al., 2018)). The same area was then first photo-bleached by a high laser intensity and then imaged with a moderate laser intensity optimized for the fluorophore lifetime lasting for 3,000 frames with an exposure time of 30ms per frame, resulting in a high-resolution image containing ~100,000 individual localizations (Figure 1B, *bottom*). The overall phase boundary did not undergo obvious change during the imaging process as the shapes and boundaries of the condensed droplets in the system remained essentially the same (Figure 1B).

Due to the stochastic nature of fluorophore switch on and off, the reconstructed super resolution image could be treated as static molecular distributions of labeled molecules in both dilute and condensed phases. Based on this super resolution image, we could define those areas that have higher localization densities as the condensed phase regions, and the rest as the dilute phase regions (Figure 1C). Accurate phase boundaries could be clearly visualized for each condensed region. Comparing the average localization densities in the condensed and dilute phases, we could estimate the partition coefficient of ~61 for NR2B (i.e., NR2B was enriched into the condense phase by ~61 folds). The calculated NR2B enrichment derived from the super-resolution imaging study was close to the value obtained by a bulk fluorescence imaging-based method shown in our previous study (Zeng et al., 2018). We noted with interest that the distribution of NR2B in the condensed phase are not homogeneous (Figure 1C), indicating formation of nanodomain-like clusters within the condensed phase.

### Simultaneous single molecule tracking in different phases

The localizations obtained from dSTORM images contained information about distributions as well as mobilities of molecules in both phases. However, the diffusion mode and densities of molecules are very different in condensed and dilute phases. A striking feature is that molecules tend to experience transiently confined state in the condensed phase (Supplemental movie 1). We developed an adaptive single molecule tracking algorithm that could automatically and robustly define optimal search ranges for molecules in different phases, and the method could effectively minimize the global assignment errors in tracking molecules in both condensed and dilute phases (Figure 2A and Figure 2—figure supplement 1-4; see “Methods” for extended description of the algorithm). Briefly, phase boundaries were determined by densities of localizations at the beginning. A default search range (500 nm) was used to assign all localizations into tracks in both condensed and dilute phases. Diffusion coefficients of NR2B in both dilute and condensed phases were estimated for determining optimized search range for different phases. All localizations were reassigned with the optimized search range to obtain final tracks of NR2B in both condensed and dilute phases.

**Figure 2:**
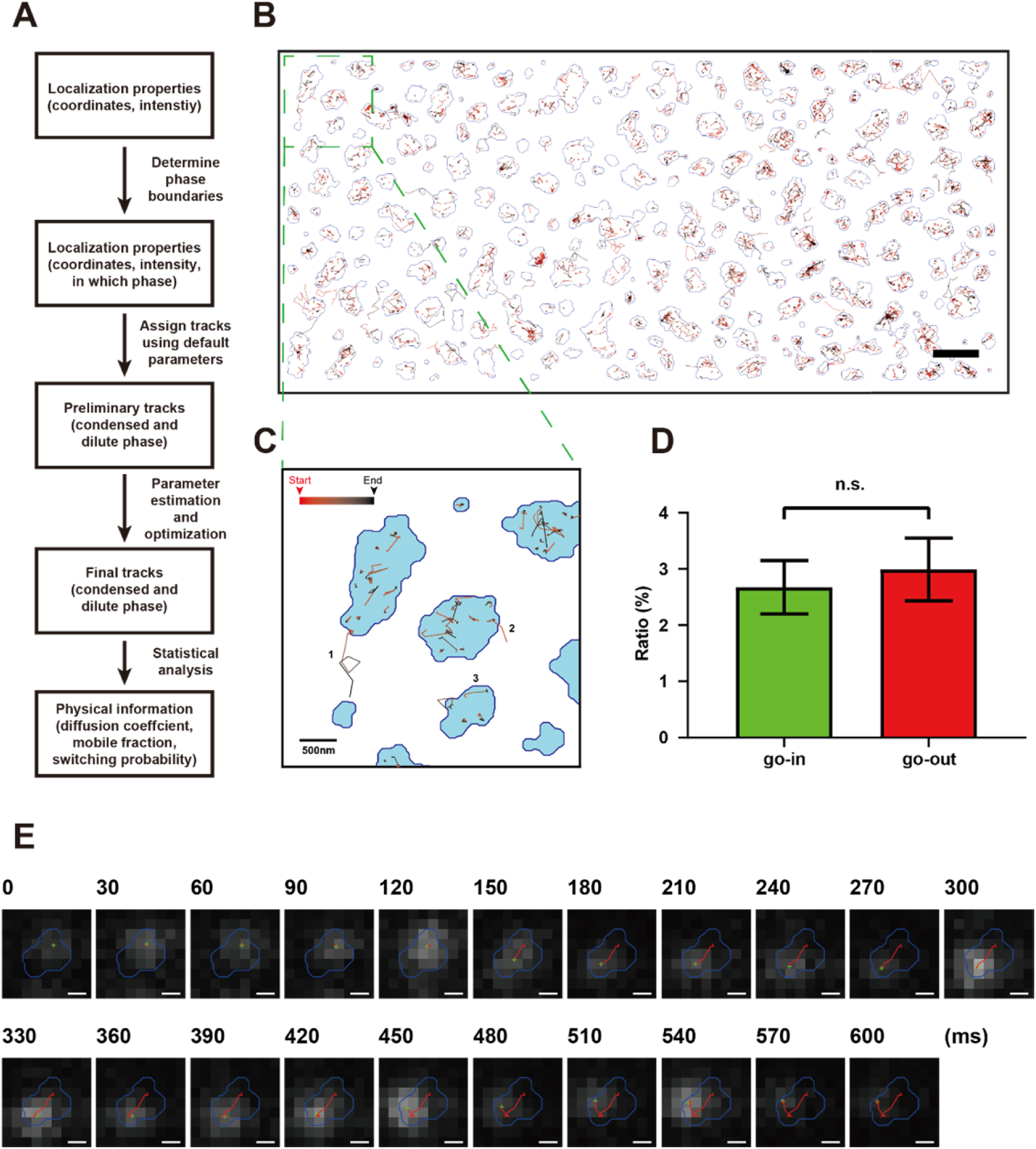
Development of an adaptive single molecule tracking algorithm for imaging single molecules in the condensed and dilute phases simultaneously. (A) Flow chart of the adaptive single molecule tracking algorithm. (B) Assignments of motion tracks of NR2B in both condensed and dilute phases in the PSD condensates formed on SLB. Each track is color coded from red to black representing from the beginning to the end of the track. The boundaries of the condensates are marked by blue lines. Scale bar: 2 μm. (C) Representative tracks showing typical NR2B motions that contains different state including exchange events of molecules between condensed and dilute phases (track 1&2), and both transiently confined and freely diffusing states in the condensed phase (track 3). (D) Percentages of NR2B molecules exchange from the dilute phase into the condensed phase (“go-in”) and exchange from the condensed phase out to the dilute phase (“go-out”) were counted in four sessions dSTORM imaging experiments. No significant difference between go-in ratio and go-out ratio was detected. P=0.378 using paired t test. (E) Raw image data superimposed with phase boundary (blue line), molecule localization (green cross) and track steps (red line) shown a typical trajectory of a molecule that experience multiple motion switchs between confined and mobile states in the condensed phase. Scale bar: 200 nm.

After adaptively assigning all localizations into tracks, we could obtain single molecule tracks of molecules in both condensed and dilute phases (Figure 2B). With this method, we could directly record events of molecules entering into and escaping from the condensed phase as well as switch motions of molecules converting between confined state and mobile state in the condensed phase (Figure 2C). The number of NR2B molecules entering into and escaping from the condensed phase were equal (Figure 2D), a finding that is consistent with the bulk equilibrium state of the PSD condensate. Interestingly, NR2B molecules within the condensed phase spent a large proportion of time in the confined state and molecules could switch between confined state and mobile state (Figure 2C&E). No confined state could be detected when only NR2B was tethered to SLB (Figure 3—figure supplement 1A). The above result indicated that NR2B in the condensed phase did not undergo homogeneous diffusion motions as one might expect. We also imaged motions of NR2B in the PSD condensates formed in 3D solution at the single molecular resolution and found that, in the condensed phase, each NR2B molecule spent a large proportion of time in the confined state (Figure 3—figure supplement 1B).

The histogram of NR2B displacement tracks in the condensed PSD phase has a dominant peak with very small displacements, corresponding to the large proportion of time of NR2B in the confined state. A small and relatively flat shoulder tailing the main peak represents the small proportion of time of NR2B in the mobile state with larger displacements (Figure 3A1). Fitting the histogram with a single population of NR2B undergoing Brownian motions could only cover the confined state peak of molecule but not the high-displacement tail of this distribution (Figure 3—figure supplement 1C1). In contrast, in the dilute phase, the overall displacements of NR2B are much larger and broader with no prominent peak with very small displacements (Figure 3A2). Even so, the displacement histogram of NR2B in the dilute PSD phase cannot be described by Brownian motions of single NR2B population (Figure 3—figure supplement 1C2), suggesting a presence of multiple populations of pre-percolated NR2B/PSD protein complexes even in the dilute phase. In contrast, in the control system with only NR2B tethered to SLB (i.e., no addition of any other PSD proteins), the histogram of NR2B displacements can be nicely fitted by a simple diffusion model (Figure 3B).

**Figure 3:**
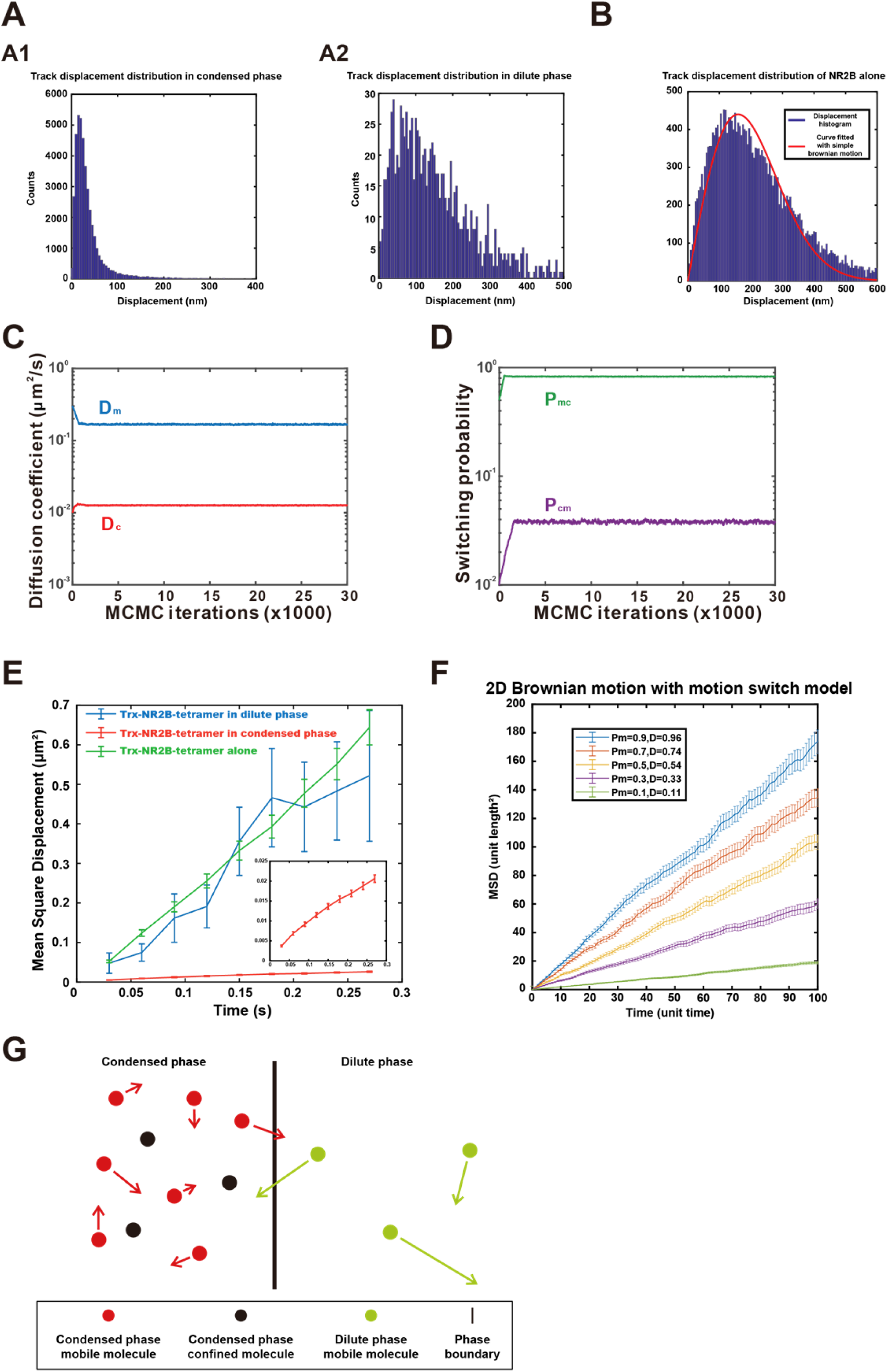
Dynamic parameters and a diffusion model for an equilibrium state phase separation system. (A) Displacement distribution of tracks in (A1) condensed phase and (A2) dilute phase. Bin size of histogram is 5 nm. (B) Displacement distribution of tracks of NR2B along tethered to SLB. Red Curve is the fitting with a simple 2D Brownian motion distribution use non-linear least squares method using MATLAB, R^2^ = 0.91, RMSE = 42.5. Bin size of histogram is 5 nm. (C&D) Fitting of the dynamic parameters of NR2B in the PSD condensates formed on SLBs with Hidden Markov Model assuming that NR2B is undergoing a two-state motion model (i.e. a transient confined state and a mobile state) in the condensed phase. The parameters are diffusion coefficient of the confined state in condensed phase (D_c_) and diffusion coefficient of the mobile state (D_m_), switching probability from the confined state to the mobile state (P_cm_) and the reversed switching probability (P_mc_). (E) Determination of the diffusion coefficients of NR2B in dilute phase (blue) and condensed phase (red) by fitting the MSDs against time with a linear regression. The figure also includes the curve and fitting of NR2B alone tethered to SLB (green). The insert shows a y-axis zoom-in view of NR2B in the condensed phase. The number of trajectories used in the fittings were 2443 for the condensed phase, 13 for the dilute phase, and 248 for NR2B alone on SLB. (F) Simulation of molecule diffusion on 2D surface with the Brownian motion with motion switch model under different mobile ratio (Pm), diffusion coefficient (D) was fitted with the MSD curve and normalized to mobile ratio = 100% scenario. (G) Schematic diagram showing molecular motions between condensed phase and dilute phase under the equilibrium state. Black and red dots represent molecules adopting confined and mobile states, respectively, in the condensed phase. Green dots represent molecules in the dilute phase. The lengths of the arrows are to indicate different mobilities of indicated molecules.

We next used the Hidden Markov Model (HMM) to fit the motions of NR2B in the condensed phase with a two-state diffusion model (Das et al., 2009; Persson et al., 2013), motivated by the clear observation of relative immobile as well as mobile NR2B molecules in the imaging experiments (Figure 2E). The parameters included the diffusion coefficients of the molecule in transiently confined and mobile states (D_c_ and D_m_) and the switching probabilities between the two states (P_mc_ and P_cm_). Maximum likelihood estimation was used to estimate the parameters iteratively, and the parameters converged quickly after several thousand iterations of optimization (Figure 3C&D). The diffusion coefficients in the mobile state and in the confined state were D_m_=0.17 μm^2^/s and D_c_=0.013 μm^2^/s, respectively. The switching probabilities were P_mc_=82.8% (mobile to confined states) per frame and P_cm_=3.8% (confined to mobile states) per frame, respectively. The confinement ratio (P_c_), defined as the percentage of time that a molecule spends in the confined state, could be calculated by these two switching probabilities as: P_c_=P_mc_/(P_mc_+P_cm_). For NR2B in the PSD condensates, P_c_=95.6%. The mobile ratio, defined as the percentage of time that a molecule spends in the mobile state, could be calculated as: P_m_=1-P_c_=4.4%. The diffusion coefficient of the molecule in each phase could be extracted by fitting the MSD (mean square displacement) values as a function of time. The diffusion coefficient of NR2B in the dilute phase of the PSD condensate is ~0.47 μm^2^/s, which is very close to that of NR2B alone tethered to SLB (~0.61 μm^2^/s) (Figure 3E). The apparent diffusion coefficient of NR2B in the condensed phase derived by fitting MSD vs time is ~0.017 μm^2^/s (Figure 3E). This fitted diffusion constant should be considered as an apparent diffusion constant (D_a_) as it contains information of the confined states and the mobile states of the molecules in the condensed phase. When the confinement ratio is large, the apparent diffusion coefficient will be dominated by the confined state and significantly differ from the diffusion coefficient in the mobile state. The simulation results of molecular diffusions of condensates formed on 2D SLB with different motion switch conditions based on free diffusion with motion switch model are consistent with the experimental trend (Figure 3F).

### A diffusion model for equilibrium liquid-liquid phase separation

We next developed a simple diffusion model to describe a phase separation system based on parameters measured using our adaptive single molecule tracking method. Consider a small region near the phase boundary (Figure 3G). The dilute phase contains sparse and fast-moving molecules. The condensed phase contains dense and slow-moving molecules which can be further categorized into either in mobile state or transiently confined state. We did not observe any obvious hinderance against motions when molecules cross the phase boundaries, thus the energy barrier at the interface between the condensed and dilute phases is likely negligible (Brangwynne et al., 2011; Feric et al., 2016). We assume that the number of molecules crossing the boundary from one side to the other are proportional to the diffusion coefficient (D) and the molecular density (σ). The influx of molecules from the dilute phase to the condensed phase can be written as J_dc_ = kσ_d_D_d_, where k is a constant, σ_d_ is molecule density in dilute phase, and D_d_ is the diffusion coefficient in dilute phase. Since a portion of molecules in the condensed phase are transiently confined, the efflux of molecules from the condensed phase to the dilute phase is J_cd_ = kP_m_σ_c_D_m_, where P_m_ is the mobile ratio of molecules in the condensed phase, σ_c_ is molecule density in condensed phase, D_m_ is the diffusion coefficient in mobile state. The net molecule flux should be zero at the equilibrium state, thus we have J_dc_ = J_cd_ or σ_d_D_d_ = P_m_σ_c_D_m_. The enrichment fold (EF) of molecules in the condensed phase over the dilute phase is defined as σ_c_/σ_d_. Thus:

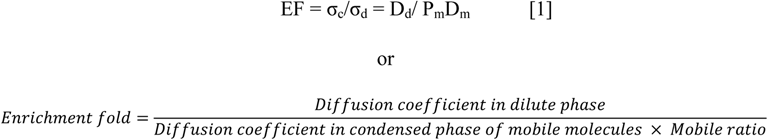

This equation connects the variables at the microscopic level (diffusion coefficient and mobile ratio) to the macroscopic level observables (enrichment fold) with a very simple relationship. If the molecules have multiple diffusion states, we can readily extend the model, viz.,

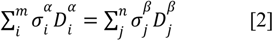

where molecules in phase α and β contains, respectively, m and n types of diffusion states. When there are multiple (two or more) coexisting phases, the sum of the products of molecule density and diffusion coefficient in different states should be equal for all the coexisting phases. The above analysis can be easily extended to phase separations in 3D solutions by simply replacing the present 2D molecule densities with corresponding 3D molecular concentrations.

Taking NR2B in the PSD condensates formed on SLB as an example, the measured diffusion parameters are: D_d_ =0.47 μm^2^/s, D_m_=0.17 μm^2^/s, and P_m_=4.4%. The theoretical enrichment fold of NR2B in the condensed phase based on *eq.* 1 is D_d_/P_m_D_m_=62.8, a value that is very close to value (~61) derived from the experimentally observed localizations (Figure 1B). The model further provides mechanistic insights into some unique properties of molecules in the condensed phase. For example, although concentrations of molecules in the condensed phase are much higher than those in the dilute phase, it is remarkable that the diffusion coefficient for the mobile fraction of molecules in the condensed phase is not dramatically different from that in the dilute phase (0.17 vs 0.47 μm^2^/s). The fraction of time that a molecule spends in the mobile state *vs* the confined state dramatically influences the macroscopic properties of the molecule in the condensed phase. For instance, NR2B spends over 95% of the time in the confined state in the PSD condensate. Accordingly, the apparent diffusion constant of NR2B in the condensed phase is small at 0.017 μm^2^/s instead of 0.17 μm^2^/s measured for the mobile NR2B fractions in the condensed phase. Additionally, the enrichment fold (or factor of enrichment) of a molecule into the condensed phase upon phase separation is also dominantly reflected by the fraction of time molecules spent in the mobile state vs that in the confined state. As we will demonstrate below, the binding affinity between molecules in the phase separation system and the molecular network complexity in the condensed phase determine the fraction of the time that a molecule spends in the mobile state vs confined state as well as the enrichment of molecules in the condensed phase.

We further validated the diffusion model with simulations of molecular motions from homogeneously mixed state towards a phase separated equilibrium state. A 2D simulation box with size of 15×30 μm^2^ and periodical boundary conditions was prepared (Figure 3—figure supplement 1D). Diffusion coefficients for the simulated molecule in the condensed phase (for mobile state only) and in the dilute phase, mobile ratio, and mobile state lifetime (derive from switching probability) of the molecule in the mobile and confined states in the condensed phase were defined as simulation input (Table 1). The Monte Carlo method was used to simulate a total of 50,000 molecules diffusing in the box for 100 seconds for each condition. The enrichment fold and the ratio of molecules exchange between condensed and dilute phases (exchange ratio) along the simulation trajectory were determined. Every simulated system eventually reached equilibrium. The enrichment fold of the molecule at the equilibrium state under each condition matched the theoretical value calculated by *eq. 1* (Table 1).

Taken together, we now have a diffusion model for the equilibrium state of a phase separation system. This model explicitly connects a set of measurable microscopic molecular motion properties with the observable macroscopic parameters of molecules in the system. The experimental method and the theoretical model developed above are simple and robust to be implemented for analyzing biomolecular phase separations in general.

### Dynamic molecular networks in condensed phases

Molecules or molecular complexes in dilute solution obey the diffusion law. In sharp contrast, NR2B molecules in the PSD condensates formed on SLB and in 3D solution spend a very large proportion of time in the immobile/confined state as if the PSD condensates can form some sort of very large and thus immobile molecular network (a process known as percolation in associative polymers including biopolymers; (Choi et al., 2020; Harmon et al., 2017; Winnik and Yekta, 1997)) capable of trapping NR2B tansiently. Since molecular processes such as molecular interactions, chemical reactions, etc. require molecules to be able to collide with each other, phase separation-mediated immobilization of biomolecules in condensed phases can have huge implications on numerous fundamental properties of these molecules (e.g., binding kinetics, catalytic speed and specificity of enzymes, spatial distributions in cellular sub-compartments, etc.).

We next asked how such network-like structure might form and what factors determine the network stability in the condensed phase of a biological condensate. We hypothesized that a phase separated system driven by multivalent and strong inter-molecular interactions, such as the PSD condensates studied above, would form highly stable and larger molecular network in the condensed phase. Accordingly, molecules bound to the network might be considered as immobile or confined by the network. In contrast, molecular networks in the condensed phase formed by weak but also multivalent molecular interactions (e.g., Intrinsic Disordered Region (IDR)-mediated phase separations) would be more dynamic and the size of the network would also be smaller (see Figure 4A for a scheme). One might envision that the molecular networks formed in the condensed phase could locally break or reform, as molecules within the network could still undergo binding and unbinding processes. Thus, the molecular networks formed in condensed phase are dynamic. The fraction of time that a molecule stays on the network is directly proportional to the binding affinity (i.e., the off-rate of the molecule from the network, a value directly related to the dissociation constant of the binding) and avidity (the available binding sites in the vicinity of the molecule, a parameter related to valency of the molecular interactions in the system) between the molecule and the network.

**Figure 4:**
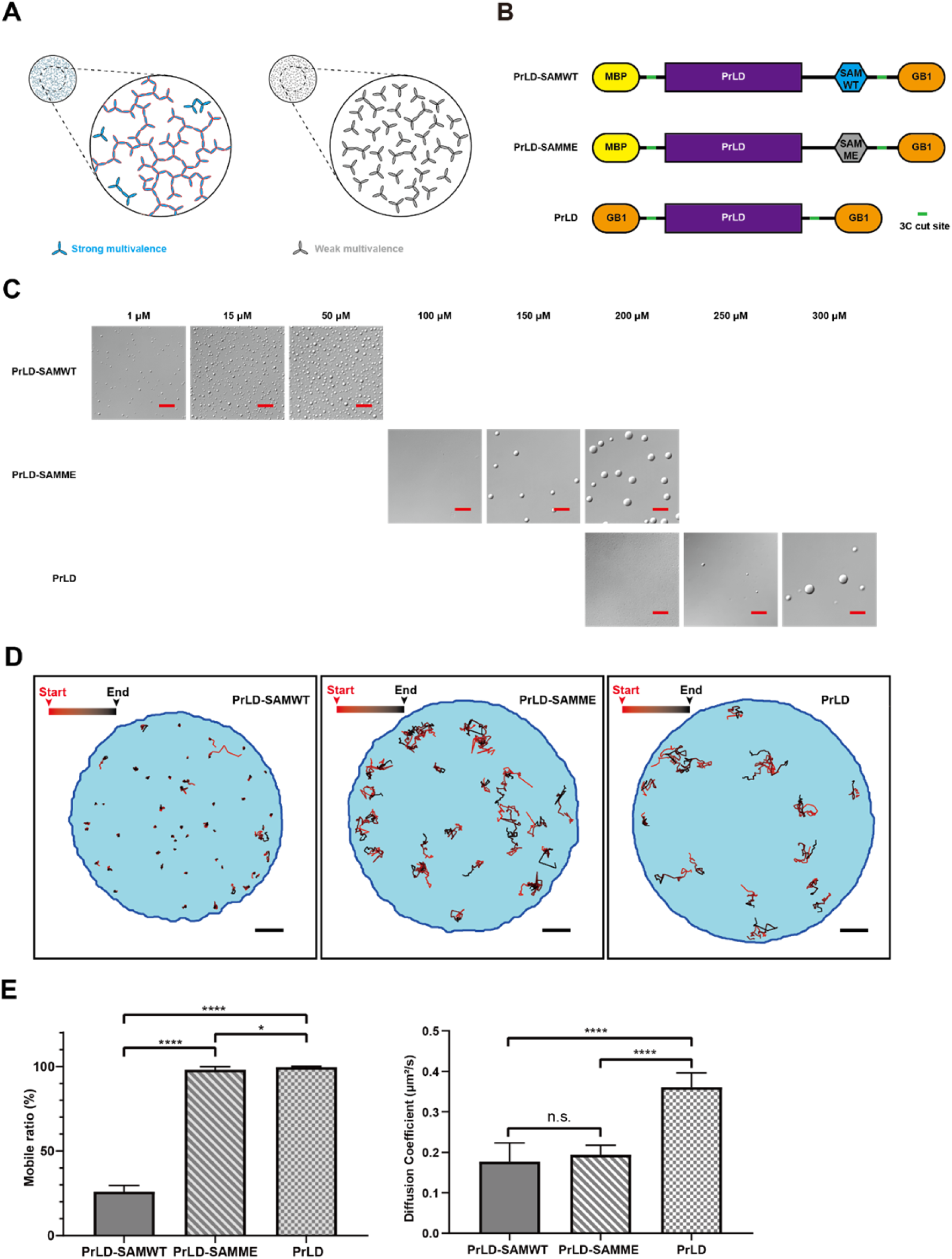
Immobilization of molecules by the large and dynamic molecular network in the condensed phase of phase separated systems. (A) Schematic diagrams showing large and stable molecular networks in the condensed phase formed by specific and multivalent interactions (left, blue) and small and dynamic molecular networks in the condensed phase formed by weak, nonspecific but multivalent interactions (right, gray). Red edge highlights the large dynamic network. (B) Schematic diagram showing composition of three designed and “caged” single protein phase separation systems with different interaction properties. PrLD, prion-like domain of FUS; SAMWT, WT SAM domain from Shank3; SAMME, the M1718E mutant of Shank3 SAM domain; MBP, maltose binding protein as a caging tag; GB1, the B1 domain of *Streptococcal* protein G as another caging tag. The HRV-3C cleavage sites (“3C cut site”) of the proteins are also indicated. (C) DIC images showing phase separations of the three designed proteins at different concentrations after removal of the caging tags by HRV-3C protease cleavage. Scale bar: 20 μm. (D) Representative tracks showing different motion properties of the three designed proteins in condensed phase. Scale bar: 2 μm. (E) Comparison of diffusion coefficient in mobile state (left) and mobile ratio in condensed phase (right) of the three designed proteins. N=12, data are expressed as mean ±SD with **** P<0.0001, * P<0.0332 by t-test.

To validate this hypothesis, we created a one-component phase separation system, a chimeric protein composed of the prion like domain (PrLD) of FUS connected with the SAM domain of Shank3 (PrLD-SAMWT, Figure 4B). The PrLD of FUS is a well-characterized IDR protein capable of phase separation by itself (Kato et al., 2012). The SAM domain of Shank3 can specifically interact with each other in a head-to-tail manner forming large polymers (Baron et al., 2006). Thus, the PrLD-SAM chimera contains a weak and multivalent interaction domain and a specific and multivalent interaction domain within one protein. The PrLD-SAM chimera could undergo phase separation at very low concentrations (see below), so we “caged” the chimera with highly soluble maltose binding protein (MBP) at its N-terminus and a small, highly soluble protein GB1 at its C-terminus (Figure 4B). The resulting “caged” protein could be purified and concentrated to as high as 500 µM. Cleavage of the caging tags of caged PrLD-SAM with HRV-3C protease induced phase separation of the protein. Substitution of Met1718 of the Shank3 SAM domain with Glu dramatically weakens the head-to-tail interaction of the domain and the mutant SAM domain (SAMME) has very weak propensity of forming oligomers in solution ((Baron et al., 2006); data not shown). We also created a “caged” PrLD-SAMME chimera to investigate the impact of weakening specific interaction on the molecular network formation in the condensed phase (Figure 4B). Lastly, we used the FUS PrLD only to investigate the property of the molecular network in the condensed phase that is solely formed by a weak and multivalent IDR sequence (Figure 4B). Again, we “caged” FUS PrLD at its both termini with GB1, so that the caged PrLD can be purified in its native form and concentrated to very high concentrations without phase separation. We predicted that the molecular networks in the condensed phase formed by PrLD-SAMWT would be the largest and most stable followed by PrLD-SAMME, and the molecular network of the PrLD condensed phase should be the most dynamic.

We compared the threshold concentrations of the three proteins for phase separation to occur. Phase separation of each protein was induced by mixing 1 µM HRV-3C protease and immediately injecting the digestion mixture into a sealed, home-made chamber incubated at 20°C. All caged proteins were completely digested within ~30 min after addition of HRV-3C protease. However, the phase separation of PrLD or the PrLD-SAMME chimera took up to 12 hours to occur (i.e., with very slow nucleation rates). Thus, we compared DIC images of the three cage-cleaved proteins captured at 12 hours after addition of HRV-3C protease (Figure 4C). The PrLD-SAMWT chimera underwent phase separation at concentration as low as 1 μM. In contrast the threshold phase separation concentrations for PrLD-SAMME and PrLD were much higher (~150 μM and ~250 μM, respectively). These results demonstrate that specific and multivalent interactions act in concert in biomolecular condensates and that the latter can dramatically lower the threshold concentration of phase separation (Espinosa et al., 2020; Lin et al., 2022; Riback et al., 2020; Zeng et al., 2018; Zeng et al., 2016).

We then compared the motion properties of the three proteins in the condensed phase use the adaptive single molecule tracking method developed above. A concentration somewhat higher than the phase separation threshold was used for each protein (i.e., 50, 200, 300 μM for PrLD-SAMWT, PrLD-SAMME, and PrLD; respectively). Each protein was very sparsely labeled with Alexa 555 (0.02% for PrLD-SAMWT, 0.005% for PrLD-SAMME, and 0.005% for PrLD) to obtain sparse but long lifetime (>1s) single molecule tracks. It is clear that, in the condensed phase, the motion properties of PrLD-SAMWT are dramatically different from those of PrLD-SAMME and PrLD (Figure 4D). Each PrLD-SAMWT molecule spent most of their time (74.0±3.7%) in the transiently confined state, but these molecules were able to switch between the confined and mobile states (Figure 4E). In contrast, the mobilities of PrLD-SAMME or PrLD in the condensed phase were much higher. PrLD-SAMME spent 98.1±1.8% of time in the mobile state and PrLD spent 99.6±0.4% in the mobile state (Figure 4E). These findings indicated that PrLD-SAMWT molecules in the condensed phase formed a very large but still dynamic molecular network due to the presence of specific and multivalent SAM-SAM interaction. In contrast, the molecular networks in the condensed phase formed solely by weak and multivalent interactions were much smaller and more dynamic. Interestingly, the diffusion coefficients of PrLD-SAMWT and PrLD-SAMME in their mobile state were very similar (0.18±0.05 μm^2^/s vs 0.19±0.02 μm^2^/s) (Figure 4E), indicating that PrLD-SAMWT and PrLD-SAMME have similar molecular size in their mobile state. The mobile state of both proteins likely corresponds to each molecule not bound to the large molecular network in the condensed phase. The diffusion coefficient of the mobile state of PrLD is 0.36±0.04 μm^2^/s (Figure 4E) and the molecular weight of PrLD is about half of PrLD-SAMWT and PrLD-SAMME, again suggesting that the mobile state of PrLD likely corresponds to the network unbound and monomeric form of PrLD. Taken together, the above single molecule tracking study revealed that strong multivalent interactions could lead to formation of large and more stable molecular networks in the condensed phase, which could dramatically reduce the overall motions of the molecules in the condensed phase. Most prominently, proteins that bind to large and stable molecular networks no longer obey the free diffusion law found in dilute solutions. Instead, in such condensed phase, proteins switch between immobile/confined state and free diffusion state, corresponding respectively to the network-bound and free forms. In the condensed phase formed by weakly interacting IDR sequences, the molecular network in the condensed phase is very dynamic and much smaller resulting in IDR proteins displaying simple diffusion-like behavior.

### Fluorescence recovery after photobleaching (FRAP) in phase separation systems

FRAP assays are widely used to examine dynamic properties of phase separation systems. Quantitative theory for analyzing FRAP results in phase separation systems based on Flory-Huggins theory have also been developed for weak interaction systems (Hubatsch et al., 2021). For heterogeneous condensed phase systems, traditional FRAP experiment might not be a good way to test the liquid-like properties (McSwiggen et al., 2019). As we have shown above, motion properties of molecules in the condensed phase are radically different for the systems involving specific multivalent interactions compared to the system with only weak interactions. Seeking a better understanding, we simulated FRAP properties of several different phase separation systems pertinent to our experiments. A 2-dimensional phase separation system mimicking phase separation on a flat surface such as lipid membranes was constructed with three different sizes of round-shaped condensed phase (radius of 0.5, 1, and 2 μm) in a box with periodic boundary (Figure 5A). By monitoring the molecule positions for 100 seconds after photobleaching, we can simulate the FRAP curve of any region in the box. We define a region with certain size to be bleached as region of interest (ROI).

**Figure 5:**
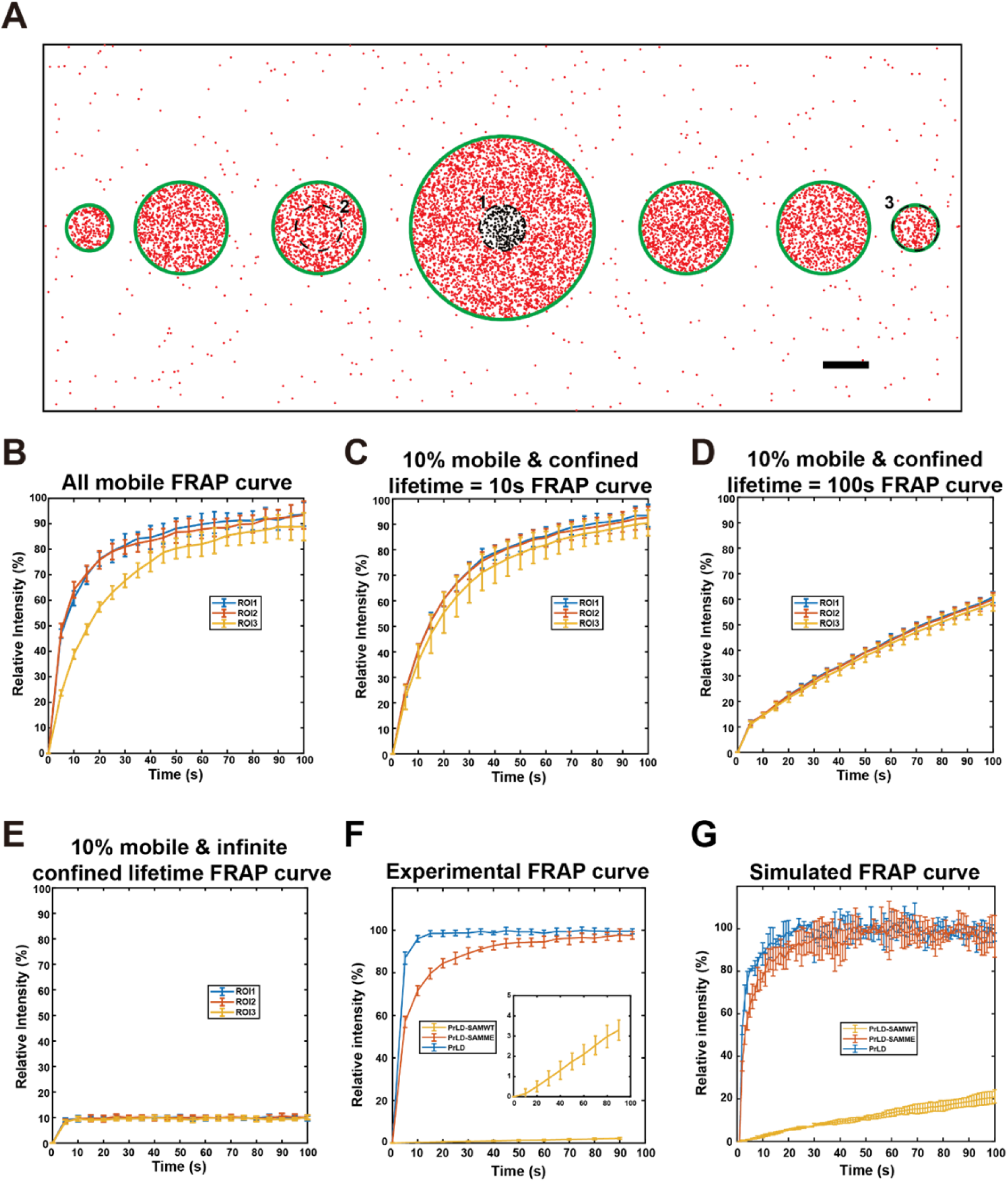
Simulations of FRAP experiments and comparison with the experimental FRAP results. (A) Schematic diagram of the phase separation system for the FRAP simulations. The simulation region is a 20 μm × 8 μm box with periodic boundary. Three ROIs (1,2,3) with a fixed diameter of 0.5 μm and positioned at the center of three different sized droplets (2, 1 and 0.5 μm in diameters, respectively) were selected for photobleaching. Green lines indicate the phase boundaries of the droplets. Black dots represent bleached molecules that can exchange with unbleached molecules in red. scale bar: 1 μm. (B) FRAP curves of the three ROIs under the condition that all molecules in the condensed phase are mobile, with an enrichment fold of 100, and diffusion coefficients in condensed and dilute phase of 0.01 μm^2^/s and 1 μm^2^/s, respectively. Data are expressed as mean ±SD from 10 repeats of simulations. (C) FRAP curves of the three ROIs under the condition that only 10% of molecules in condensed phase are mobile and diffusion coefficients of the molecule in the condensed and dilute phase of 0.1 μm^2^/s and 1 μm^2^/s, respectively. The lifetime of the molecule in the confined state was set at 10 seconds. (D) Same as in *C* except that the lifetime of the molecule in the confined state was set at 100 seconds. (E) Same as in *C* except that the molecule in the confined state were treated as permanently immobilized. (F) Experimental FRAP curves of PrLD, PrLD-SAMME, and PrLD-SAMWT condensates. In each case, a photobleaching region with the size of 1.95 μm in diameter was selected inside a large droplet (see Figure 5—figure supplement 1). The zoom-in panel is an expanded view of the FRAP curve of PrLD-SAMWT. Data are expressed as mean ±SD, with recovery experiments performed on 10 different droplets. (G) Simulated FRAP curves of the three designed proteins in the condensed phase using the parameters derived from the experiments described in Figure 4. The region selected for photobleaching is with a diameter of 2 μm and located in condensed phase with infinite size. Data are expressed as mean ±SD from three simulations.

We first asked whether the size of a ROI vs the size of a condensed phase may affect FRAP results. For this simulation, the diffusion coefficients of the molecule were set at 0.01 μm^2^/s and 1 μm^2^/s for condensed and dilute phases, respectively. The mobile ratio was set at 100% (i.e., simulating FRAP curves for a phase separation system dominated by weak multivalent interactions). This setting leads to 100 times enrichment of the molecule into the condensed phase. The ROIs with a radius of 0.5 μm were selected at the center of the three different sized droplets (ROIs 1~3, Figure 5A). The simulated results showed that the apparent recovery curve for ROI3 was considerably slower than the first two ROIs (Figure 5B), as all molecules in ROI3 needed to diffuse from the dilute phase into the entirely bleached condensed phase. This simulation indicates that one should select condensed phases with similar sizes when comparing FRAP curves of related phase separation systems. When possible, always select a small ROI within a large droplet for FRAP analysis.

We next simulated FRAP curves for the phase separation systems containing specific interactions. We first set the mobile ratio of simulated molecules in condensed phase at 10%, with average lifetime of the confined state being 10 seconds. To maintain the same apparent diffusion coefficient and the same enrichment level in the condensed phase as those in the system with no molecular confinement described above, the diffusion coefficient of the molecule in the mobile state within the condensed phase was set as 0.1 μm^2^/s. We simulated the FRAP curves for the same three ROIs as indicated in figure 5A. Although the FRAP recovery speed for ROI3 was still slower than the other two ROIs, the difference became smaller (Figure 5C). Further increasing the lifetime of the confined state to 100 seconds while keeping other parameters unchanged led to linear-like slow recovery curves and the differences among the three ROIs were further diminished (Figure 5D). In an extreme case where the lifetime of the confined molecules were infinitely long (i.e., extremely strong binding and resulting in behaviors similar to those of unrecoverable aggregates), all three recover curves looked similar with a fast recovery speed to their maximal recovery level at 10% (Figure 5E). These simulations suggest that the property of the molecular network in the condensed phase can have dramatic impact on its FRAP recovery rate in the system. For example, a slow recovery speed from a FRAP experiment does not necessarily mean that the molecule in condensed phase moves slowly. Instead, it may mean that the molecule spends most of the time bound to the molecular network. Once switched into mobile state, it can diffuse quite freely and rapidly.

We validated our simulations by experimentally measuring FRAP curves of the three phase separation systems shown in Figure 4B. A small ROI with identical radius within a relatively large condensed droplet was selected for the FRAP experiments in each of the three systems (Figure 5—figure supplement 1). The PrLD system showed the fastest recovery speed, and the PrLD-SAMME system showed a slightly slower recover speed. The PrLD-SAMWT system, with its large and stable molecular network in the condensed phase, displayed a very slow and near linear recovery curve (Figure 5F). We further simulated the FRAP curves of the three systems using the diffusion coefficients and confinement ratios derived from the single molecule tracking data (Figure 4E). The simulated FRAP curves were overall very similar to those obtained experimentally (Figure 5G vs 5F). The faster recovery rate in the simulation curve vs the experimental curve for the PrLD-SAMWT system is likely due to the overestimations of the mobile ratio of the protein in the tracking experiment. Taken together, the above theoretical and experimental studies revealed that the FRAP curve of a molecule in a phase separation system is heavily influenced by the proportion of time of the molecule transiently trapped in the confined state, a parameter that is directly linked to the specific multivalent interactions of the system. A low recovery rate in the FRAP assay does not necessary mean that a large fraction of molecules in the condensed phase is permanently immobile.

## Discussion

In this study, we developed a method that can track single-molecule motions in both condensed phase and dilute phase simultaneously by using photo-switchable dye labelled proteins. To accommodate the heterogeneity of both the distributions and diffusion modes of molecules in the condensed and dilute phase, an adaptive single-molecule tracking algorithm was developed by setting the optimized search range for molecules with different diffusion coefficients. The method is simple and highly robust. It can be deployed to track single-molecule motions of phase separation systems with very broad dynamic ranges including highly dynamic systems formed by IDR proteins or very stable (or highly percolated) systems formed by strong and specific multivalent molecular assemblies such as PSDs. With implementations of sparse labeling techniques such as HaloTag, our method can be applied to track single-molecule motions of biological condensates in living cells.

The most important finding from our study is that molecules in the condensed phase do not exhibit simple diffusion behaviors as those observed in dilute solutions. Instead, molecules constantly switch between transient confined state and mobile state in the condensed phase, a phenomenon that is most likely underpinned by phase separation-mediated formation of large molecular networks. The size and dynamic properties of the molecular network in the condensed phase are determined by the binding affinity (or affinities) and valence of the interaction(s) of the molecule(s) in the phase separation system, a finding that has been recently predicted by theoretical simulations treating biomolecules are associative polymers (Choi et al., 2020; Harmon et al., 2017). The fraction of time and the time duration that a molecule spends at the confined state is also determined by the binding affinity of the molecule to and the dynamic property of the network. Surprisingly, the diffusion coefficients of mobile-state molecules in the condensed phase are only slightly lower than the counterpart molecules in the dilute phase. For biological condensates of which their formations are largely driven by specific molecular interactions (it is our opinion that most cellular condensates belong to such category; (Chen et al., 2020; Feng et al., 2021)), molecules in the condensed phase spend most of their time in the confined state. Since the fraction of time and the time duration that a molecule spends in the confined state vs those in the mobile state are basic defining parameters for the functions of molecules in any reaction systems (e.g., binding/unbinding rates, kinetics of enzyme catalysis, lifetime/dwelling time a molecule in a molecular machinery, etc.), our finding reveals a fundamental aspect of molecular properties created by biological condensates that is distinctly different from that in dilute solutions. Our study implies further that, in theory, unlimited types of biological condensates with very broad dynamic network properties may form using the existing repertoires of proteins and nucleic acids via different combinations of binding affinities and interaction valences. Thus, phase separation-mediated formation of biological condensates is a very powerful means for cells to form numerous subcellular organelles with a continuum dynamic and material properties ranging from very dynamic, dilute solution-like assemblies to highly stable, solid-like systems. Confinements of molecules in cellular condensates have been observed by single-molecule tracking experiments recently (Chong et al., 2022; Miné-Hattab et al., 2021; Niewidok et al., 2018). However, since the molecular compositions of cellular condensates cannot be easily defined, the mechanistic bases underlying the confined state of molecules in these cellular condensates are difficult to be discerned. Single molecule tracking experiments using the reconstituted and compositionally defined phase separation systems in the current study allowed delineation of the mechanism underlying the unique motion properties of molecules in the condensed state.

During our adaptive single molecule imaging process, we did not observe obvious motion speed slow down or enhancement when molecules enter or leaving a condensed phase in all condensate systems investigated in this work. Therefore, the tension (or depth of the energy well) at the interface between the condensed phase and the dilute phase in each condensate is not large enough to significantly impact molecular exchanges between the two phases.

Since the interactions between molecules can be modulated via numerous cellular processes such as posttranslational modifications, protein biogenesis/turnovers, epigenetic modification, cellular milieu alterations, etc., the dynamic network properties and consequently the functions of organelles formed via phase separation may be regulated in ways that are distinctly different from those occurring in dilute solutions. Compared to the rich knowledge to and quantitative theories for the dilute-solution systems, few satisfying theoretical frameworks have been established for the condensed assemblies formed via phase separation in cells (see (Mittag and Pappu, 2022) and refs therein), partly due to our poor understandings of microscopic motion properties of molecules in the condensed phase. The dramatic dynamic and material property differences of condensates formed by weakly associative IDR proteins and by biomolecules with specific interactions indicate that biological phase separation research has only touched the tip of the iceberg, given that the vast majority of research only deals with IDR proteins.

## Materials and Methods

### Protein expression and purification

Constructs for expression of Trx-His-GCN4-NR2B, PSD-95 (UniProt: P78352-1), Shank3 (UniProt: Q4ACU6), GKAP (UniProt: Q4ACU6-1), Homer3 (UniProt: Q9NSC5-1) were described previously (Zeng et al., 2018). MBP-His_8_-GCN4-NR2B tetramer was created by inserting GCN4-NR2B sequence into an in-house modified pET32a vector. MBP-His_8_-GCN4-NR2B trimer and dimer were mutated from the tetramer version by changing the hydrophobic residues in the GCN4 domain (Delano and Brünger, 1994). Constructs of GB1-PrLD-GB1 contained FUS-PrLD (UniPort: P56959, segment: 1-212) with the protecting GB1 protein fused to the N- and C-terminal ends. MBP-PrLD-SAMWT-GB1 is a fusion protein with the SAM domain of Shank3 (aa 1654-1730) fused to the C-terminal end of PrLD, and the resulting chimeric protein was further protected by tagging its N-terminus with MBP and C-terminus with GB1. MBP-PrLD-SAMME-GB1 is the same as MBP-PrLD-SAMWT except that Met1718 in the SAM domain was replaced by Glu. An additional cysteine was inserted at the N-terminus of PrLD for cysteine labeling. All constructs were confirmed by DNA sequencing. Recombinant proteins were expressed in *Escherichia coli* BL21 (DE3) cells in LB medium at 16 °C. Protein expressions were induced by adding 0.25 mM IPTG when OD_600_ reached 0.6-0.8. His_8_-tag containing recombinant proteins were purified using Ni^2+^-NTA agarose affinity column followed by size exclusion chromatography (Superdex 200 26/60 column from GE healthcare) in a final buffer containing 100 mM NaCl, 50 mM Tris-HCl (pH 7.8), 1 mM DTT and 1 mM EDTA. The purified proteins (except of the NR2B proteins for lipid binding and PrLD/PrLD-SAMME/PrLD-SAMWT) were then subject to tag removal by HRV-3C or TEV protease at 4 °C overnight followed by another round of size exclusion chromatography. All purified proteins were checked for free of nucleic acid contamination.

### Protein fluorescence labeling

His_8_-tagged NR2B proteins were labelled with Alexa 647 NHS ester (Thermo Fisher) and PrLD-SAMWT/PrLD-SAMME/PrLD proteins labelled with Alexa 555 maleimide (Thermo Fisher). Alexa 647 NHS ester was first dissolved in DMSO at a concentration of 10 mg/mL. Before labeling, all purified proteins were exchanged to a Tris-free buffer containing 100 mM NaHCO3 (pH 8.4), 100 mM NaCl, and 1 mM EDTA (plus 1 mM DTT for NR2B) using a HiTrap desalting column. NR2B was concentrated to 20-50 μM and mixed with the corresponding dye at a 1:1 molar ratio. Alexa 555 maleimide was dissolved in DMSO at a concentration of 10 mg/mL and mixed with PrLD-SAMWT/PrLD-SAMME/PrLD (>100 μM, without DTT) at a 1:1 molar ratio. The mixture was incubated at room temperature for about 1 hour and the reaction was terminated by adding 200 mM Tris-HCl (pH 8.2). The mixture was next loaded to a HiTrap desalting column to separate the unreacted fluorophores and to exchange proteins to buffers for following experiments. Efficiency of individual labelling was measured by Nanodrop 2000 (ThermoFisher). Unlabeled protein was mixed with each labelled protein to adjust the final labelling ratio needed for imaging experiments.

### Fast protein liquid chromatography coupled with static light scattering (FPLC-SLS) assay

The analysis was performed on an AKTA FPLC system (GE Healthcare) coupled with a static light scattering detector (miniDawn,Wyatt) and a differential refractive index detector (Optilab, Wyatt). Protein samples (concentrations for each reaction were indicated in the figure legends) were filtered and loaded into a Superose 12 10/300 GL column pre-equilibrated by a column buffer composed of 50 mM Tris, pH 8.2, 100 mM NaCl, 1 mM EDTA, and 2 mM DTT. Data were analyzed with ASTRA6 (Wyatt).

### Lipid preparation

POPC (Avanti lipids, Cat No:850457P), DGS-NTA-Ni^2+^ (Avanti lipids, Cat No:790404P) and PEG-5000 PE (Avanti lipids, Cat No:880230P) were first solubilized in chloroform to a stock concentration of 20 mg/mL, 10 mg/mL and 1 mg/mL, respectively. Lipid mixture containing 98% POPC, 2% DGS-NTA-Ni^2+^ and 0.1% PEG-5000 PE was dried under a stream of nitrogen gas followed by a vacuum pumping to evaporate chloroform thoroughly. The dried lipids were then resuspended in PBS to a final concentration of 0.5 mg/mL. Multi-lamellar vesicle solution was next solubilized by 1% w/v sodium cholate and loaded onto a desalting column. During the desalting process, sodium cholate will be diluted allowing small uni-lamellar vesicles (SUVs) to form in the buffer containing 100 mM NaCl, 50 mM Tris-HCl (pH 7.8), and 1 mM TCEP (the 2D buffer).

### Coating chambered cover glass with lipids

Chambered cover glass (Lab-tek) was washed with Hellmanex II (Hëlma Analytics) overnight, thoroughly rinsed with MilliQ H_2_O. The chambered cover glass was then washed with 5 M NaOH for 1 hr at 50°C and then thoroughly rinsed with MilliQ H_2_O. The cleaned coverslips were washed three times with the coating buffer (50 mM Tris, pH 8.2, 100 mM NaCl, 1 mM TCEP). Typically, 150 μL SUVs were added to a cleaned chamber and incubated for 1 hr at 42 °C, the SUVs would fully collapse on glass and fuse to form supported lipid bilayers (SLBs). Chambers with SLBs were then gently washed three times each with 750 µl of coating buffer to remove extra SUVs before being blocked by the clustering buffer (coating buffer plus 1 mg/ml of BSA) for 30 mins at room temperature.

### Phase separation on SLB

The supported lipid bilayers contained 2% DGS lipid with Ni^2+^-NTA attached to its head. We used GCN4-NR2B with an N-terminal thioredoxin (TRX)-His_8_ tag (referred to as NR2B tetramer) to attach to SLBs via binding to DGS-NTA-Ni^2+^. The NR2B (4µM final concentration) tetramer was added to a SLB-containing chamber. After 30 mins incubation at room temperature, the chamber was washed with the clustering buffer for three times (each time at 750 µl volume) to remove excessive NR2B tetramers. PSD-95, Shank3, GKAP and Homer3 (each at 2 µM final concentration) were sequentially added into the system. Imaging acquisition started at 15 mins after adding all components.

### Phase separation in 3-dimensional solution in chamber

For PSD condensates, 10 µM of 5 PSD proteins were mixed and injected into a homemade chamber and sealed immediately (Zeng et al., 2016). The mixtures were incubated for 15 mins before starting image acquisitions. For the PrLD/PrLD-SAMME/PrLD-SAMWT systems, 300/200/50 µM of “caging” tag-containing protein was mixed with 1 µM of HRV-3C protease, each mixture was injected into a homemade chamber and sealed immediately. The samples were incubated at 20°C for 12 hrs before image acquisitions.

### dSTORM imaging

Freshly prepared imaging buffer (the 2D buffer plus 1% D-Glucose (Sigma G8270), 5.6 μg/mL glucose oxidase (Sigma G2133-50KU, from 100 x stock prepared in the coating buffer), 40 μg/mL catalase (Sigma C9322-10G, from 100 x stock prepared in the coating buffer) and 15 mM β-mercaptoethanol) was injected into an imaging glass chamber to replace the original coating buffer. Imaging of each sample was completed within 30 min upon addition of the imaging buffer.

dSTORM images for the condensates formed on SLBs were taken by a home-built two-color super-resoution localization microscope based on a Nikon Ti-E inverted microscope body (Zhao et al., 2015). Here only one channel was used to image samples labelled with Alexa 647. A 100x objective lens (CFI Apo TIRFM 100x Oil, N.A. 1.49, Nikon) was used to observe the fluorescence signals. An EMCCD (electron-multiplying charge-coupled device, Andor, IXon-Ultra) was applied to collect the emission lights that passed through a channel splitter. For each sample, 2000 frames of images with an exposure time of 30 ms/frame were captured from at least 6 different areas. The laser intensity was fixed at 1 kW/cm^2^ during the imaging and the microscope was at the TIRF mode. If single molecule signal density of a sample was too high, a pre-image photobleaching with a strong laser intensity (4.0 kW/cm^2^) was used to reduce the single molecule signal density. The TIRF raw images were processed by Rohdea (Nanobioimaging Ltd., Hong Kong) to generate the localization coordinates in each frame.

dSTORM images for 3D phase separation system were taken by a Zeiss Elyra7 microscope with a 63x oil objective lens. Samples were first bleached with a full power laser (500 mW) and then imaged with 20% of the full power of the 488/561/641 nm lasers with the HILO mode illumination. A TIRF-hp filter was used during imaging. For each sample, 4000 images were captured an exposure time of 30 ms/frame. Autofocus with the “definite focus” strategy was performed at every 500 frames. Maximum point spread function size was set at 9 and signal-to-noise ratio was set at 5 when capturing single molecules with Zeiss Elyra7. Samples were labeled with 0.005%~0.1% ratio of dyes depending on the signal density.

### Adaptive single molecule tracking algorithm

The heterogeneity of molecule distributions and diffusion modes in the dilute and condensed phases of liquid-liquid phase separation systems requires an adaptive single molecule tracking algorithm to minimize the track assign error locally and globally. Traditional single molecule tracking in high density systems (Jaqaman et al., 2008; Manley et al., 2008; Tinevez et al., 2017) usually set a global search range (step limit) manually to connect molecular tracks. A step limit that is too small will cause lots of missing connection for molecules with fast diffusions; whereas a step limit that is too large will cause lots of false positive connections for molecules with slow diffusions. Such errors cannot be fixed in post-tracking data analysis. A biological condensate system typically contains a condensed phase with slow diffusing molecules coexisting with a dilute phase with fast diffusing molecules, and molecules in the condensed phase constantly switch between mobile state and confined state. A single global maximum step limit without any prior knowledge might not suit for tracking molecules in a phase separation system. A typical solution for motion switch is to use the Hidden Markov Model (HMM) to fit a diffusion model that contains diffusion coefficients (D) and switching probabilities (P) for different diffusion states (S) (Das et al., 2009; Persson et al., 2013). Taking all above factors into consideration, we developed a new algorithm by adaptively choosing maximum step limit for different diffusion states to link the localizations into tracks and using HMM to fit a two-state diffusion model in the condensed phase.

The track assignment errors can be divided into two parts, true negatives and false positives. A true negative error is defined as an existing track was not linked, which leads to miss of long-distance steps. This part of the error can be estimated by the Boltzmann distribution if we assume that molecules undergo Brownian motion in the mobile state, 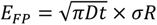 *E*_TN_ = *e*^-R2/^4Dt. A false positive error is defined as linking of a non-existing track. This part of the error can be estimated with collision frequency of particles that have a diameter same as the search range, [inlne]. Thus, the estimated assignment error under a certain search range R is:

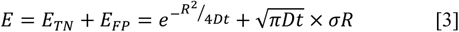

To find the minimized error of the search range R under certain diffusion coefficient D and molecule density *σ*, we just needed to find the point that 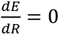 and 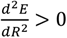. Noted that the root mean square displacement (RMSD) was 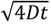, we replace 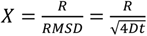 as the ratio of search range to RMSD of the molecules. The first derivative 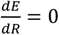 could be transformed into 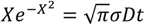. The fluorophore density in the condensed phase and the dilute phase were typically 0.20~0.40 /μm^2^ and 0.01~0.02 /μm^2^, respectively; diffusion coefficients were 0.02~0.2 μm^2^/s and 0.5~2 μm^2^/s, respectively. The solution of X under such conditions was within a very small region of 2.5~3 (Figure 2—figure supplement 1A). We could roughly estimate the diffusion coefficients by displacement distributions (Figure 3A&B) (Hansen et al., 2018) starting with a default search range (500 nm). We then moved on to find an optimized search range and use this optimized search range to complete the final track assignments (illustrated in Figure 2A).

We next validated this optimized R (search range) with simulated molecular systems undergoing homogeneous Brownian motions with different diffusion coefficients and densities. For systems that are similar to our 2D PSD system showed the same converged optimal X at ~2.5 (Figure 2—figure supplement 1B~E). We simulated many other conditions with different molecular densities (N) and moving speeds (RMSD) in both dilute and condensed phases, and all showed a similar optimal X of ~2.5 (Figure 1—figure supplement 2&3). For systems with molecules undergoing switching between confined state and mobile state in the condensed phase (defined as fraction of mobile state or mobile ratio, Pm) and with different lifetime in the mobile state tm, the optimal X value also converged to ~2.5 (Figure 1—figure supplement 4). These simulation results indicated that the optimal X value was very similar under different conditions. Thus, we used a default X=2.5 to determine the optimized R (search range) for different experiments.

To valid the algorithm experimentally, we prepared His_6_-tagged MBP fused with GCN4 dimer, trimer, or tetramer, respectively. The three fusion proteins have a molecular weights ratio of 2:3:4 measured by light scattering experiment (Figure 1—figure supplement 1F). We measured the diffusion coefficients of the three MBP proteins coated onto SLBs with our adaptive single molecule tracking algorithm without any pre-set parameters (Figure 1—figure supplement 1G). The measured diffusion coefficients for the MBP-GCN4 dimer, trimer, and tetramer are very close to 1/2:1/3:1/4, which is the expected theoretical value for the three proteins on SLB. Thus, this experimental data showed that our developed algorithm is robust in adaptively determine the diffusion coefficients without any prior knowledge.

### Generating simulated homogenous and heterogenous molecular systems

Molecules distributed homogeneously both in condensed and dilute phases were generated based on the Monte Carlo method to simulate localizations obtained in single molecule tracking experiments. Diffusion coefficients D, molecule densities N were set for different scenarios. Averaged lifetime was set to 3.5 frames based on our experimental average track length and with a Poisson distribution. Number of total tracks were calculated before simulation and every track started with a random frame following uniform distribution. For one single track with diffusion coefficient D, the displacement in a short time interval t (0.0001 s) will be:

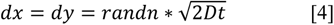

where *dx* and *dy* are the displacements along horizontal and vertical axes, *randn* is a random number with the standard normalized distribution. All data were saved into two versions, one formatted including track information as ground truth and the other formatted by frames and localizations in each frame without track information as simulated data for evaluate tracking algorithm.

Molecules distributed heterogeneously in condensed phases were generated with similar process with an additional set of switching probabilities between mobile and confined states. Molecules in a confined state will be restricted to a certain position, thus the simulated localizations will follow a Gaussian distribution based on the point spread function. A confined molecule can switch to mobile state in next frame with a probability P_cm_, and a mobile molecule can switch to a confined state in next frame with a probability P_mc_. Mobile ratio (Pm) can be calculated from the switching probability Pm, which is defined as Pm= P_cm_/(P_cm_ + P_mc_). The lifetime of the mobile state can be expressed as tm=1/P_mc_. The data were saved into two versions same as that described for the homogenous system.

Each scenario was simulated for 500-10,000 frames dependent on the molecule density, which led to a similar total track number for each system. The script for the simulations was in-house coded by MATLAB.

### Evaluation of simulated data

For each condition, simulated data were fed to the algorithm to assign localizations into tracks with different maximum step limit. According to equation 3, the ratio of maximum step limit to root mean square displacement (R/RMSD, defined as X) is the key variable, so we covered X from 1 to 5 with a step size of 0.5. Track assignment error composed of “True Negative tracks” (TN, two localizations belong to a same track in ground truth but not recognized as the same track) and “False Positive tracks” (FP, two localizations do not belong to a same track in ground truth but recognized as a same track) and calculated as the ratio of the absolute value of (the algorithm output - ground truth)/ground truth. Diffusion coefficient error was calculated by fitting mean square displacement (for the homogenous system) or fitting a two-state model (for the heterogenous system) and compared with the original setting for each condition. The scripts for the evaluations were coded by MATLAB.

### Brownian motion with motion switch model simulation

Consider a particle undergoing Brownian motion on a 2-dimensional surfacewith the additional feature that it can switch between confined and mobile states at each time step with certain switching probabilities. Specifically, during each time step, if the particle is in a confined state, it will have a probability of P_cc_=0.9 to remain in the confined state and therefore a chance of P_cm_=0.1 to switch to the mobile state. If the particle is in the mobile state, it moves in a random direction with a random displacement (i.e., following the standard normal distribution) and also have a chance of P_mc_ = P_cm_*P_c_/P_m_ to switch to confined state. To maintain a steady-state confined-mobile balance, the total switch events from mobile state to confined state and vice versa are identical, i.e., P_m_*P_mc_ = P_c_*P_cm_. Since P_c_+P_m_=1, the chance of switching from mobile state to confined state is P_mc_ = P_cm_*(1-P_m_)/P_m_. n=500 particles and simulation time length T=100 sec are used in each simulation. Mobile ratio Pm was studied from 0 to 0.9 with a step size of 0.1.

### Equilibrium state phase simulation

The phase boundaries were constructed in accordance with experiments (Figure 1B). The boundaries were not change during the simulation. An area of 15×30 μm^2^ with periodic boundary conditions was used for the simulation (Figure 3—figure supplement 1D). At the beginning of the simulation, a large number of molecules (n=50,000) were randomly distributed in the condensed and dilute phases with initial enrichment fold of 60. All molecules in dilute phase were treated as mobile during the simulation, the diffusion coefficient was set at 0.6 μm^2^/s. The motions of molecules in condensed phase consisted of those in the confined state and the mobile state. Molecules in the confined state were treated as immobile with fixed positions. Molecules in the mobile state were undergoing Brownian motion and the diffusion coefficient was set at 0.1 μm^2^/s. The switch between confined state and mobile state was determined by the switching probability P_cm_ and P_mc_. P_mc_ could be directly calculated through the averaged dwell time of mobile state (0.1 second) as the lifetime of the fluorophore (~1 second) was much longer than the molecule’s dwell time. P_cm_ was calculated using the equilibrium condition between the mobile to immobile state by:

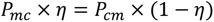

where η is the mobile ratio (10%). The simulation time step was t 0.0001 second with the total simulation time being 100 seconds. The script for the simulation was in-house coded by MATLAB.

### Fluorescence recovery after photobleaching (FRAP) assay

Proteins were labelled with Alexa Flour 555 (Thermo Fisher) at 1% for PrLD-SAMME and PrLD-SAMWT) or 0.5% (for PrLD). FRAP assays were performed on a Zeiss LSM 880 confocal microscope at room temperature. A region for bleaching (R1) with a diameter of 2 μm was selected within a large, condensed droplet. A reference region (R2) with the same size of R1 was selected in another large, condensed droplet as the system control. R1 was bleached with 40/30/10 iterations with 100% 561 nm laser power and followed by recording fluorescence intensity of the selected regions for 100 seconds in the time-lapse mode with a 10-second gap between each point for the PrLD-SAMWT system and 5 seconds for PrLD-SAMME and PrLD systems. The fluorescence intensities were normalized to 0% right after photobleaching and to 100% before photobleaching. Each data point was calibrated by recorded fluctuation of the intensity of the reference region R2.

### FRAP simulation

FRAP simulations were based on the simulation for the equilibrium state phase separation. A 20x8 μm^2^ box with periodic boundary containing seven spherical condensed droplets was used (Figure 5A). The radius and centre coordinates of condensed droplets were 0.5/1/1/2/1/1/0.5 μm and 1/3/5/10/15/17/19 μm from the left edge of the box, respectively. Conditions 1/2/3 (see Figure 5A) had the same bleaching size with a diameter of 1 μm and centered in the large/median/small droplets. The enrichment fold was set at 100 for all simulations, the mobile ratio was set at 100% or 10%, and the confined state lifetime was set at 10 or 100 seconds. For totally immobilized molecules, the lifetime of the confined state was infinite and the switching probability between mobile and immobile was zero. All simulations were carried out for 100 seconds with a time step of 0.0001 second. A total of 50,000 molecules were used for each simulation.

## Acknowledgments

This work was supported by grants from the Minister of Science and Technology of China (2019YFA0508402), National Natural Science Foundation of China (82188101), Shenzhen Bay Laboratory (S201101002), RGC of Hong Kong (AoE-M09-12, 16104518 and 16101419), and a HFSP Research Grant (RGP0020/2019) to MZ. The research effort in HSC’s group was supported by Canadian Institutes of Health Research grant PJT-155930 and Natural Sciences and Engineering Research Council of Canada grant RGPIN-2018-04351.

## Author contribution

ZS and MZ conceived the idea and designed experiments; ZS, BJ, YX performed experiments; ZS, JW, and TP performed simulations; ZS, HSC, SD and MZ analyzed data; SD supervised imaging experiments, ZS and MZ wrote and revised the manuscript and all authors provided input, MZ coordinated the study.

## Competing interest claim

The authors declare no competing financial interests.

## Supplemental Figures and Legends

**Figure 1—figure supplement 1:**
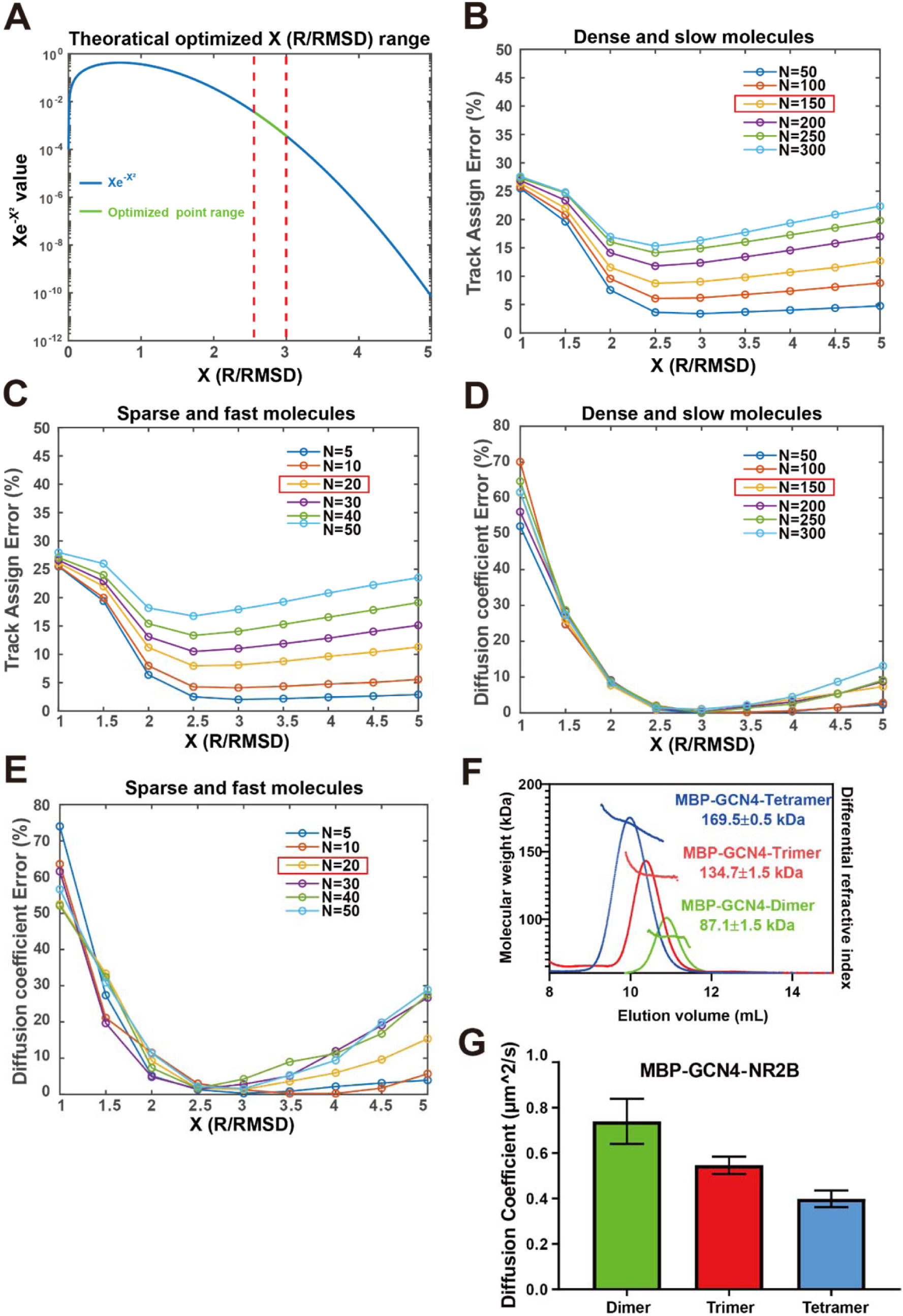
Evaluation of the adaptive single molecule tracking algorithm by simulation and by experiments. (A) Optimization point demonstration. Blue line showing the curve of left part of the equation 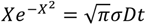, green line showing the right part range of the equation in typical scenarios of phase separation. The red dashed line showing the optimized point range in different scenarios. (B&C) Simulated track assignment errors of molecules in homogeneous condensed phase with a slow diffusion (D~0.1 μm^2^/s, panel B) or in dilute phase with fast diffusions (D~1.0 μm^2^/s, panel C) under different molecule densities (N) and maximum step limit/RMSD (R/RMSD) ratios. The red box highlighted the situation matching our experimental data for the PSD system. (D&E) Simulated diffusion coefficient errors of molecules in homogeneous condensed phase with slow diffusions (D~0.1 μm^2^/s, panel D) or in dilute phase with fast diffusions (D~1.0 μm^2^/s, panel E) under different molecule densities and R/RMSD values. The red box highlighted the situation matching our experimental data. (F) FPLC-coupled with static light scattering analysis showing the column behavior and measured molecular weight of the purified MBP-His_6_-GCN4-Dimer, Trimer, and Tetramer. (G) Diffusion coefficient of homogeneous solutions of MBP-His_6_-GCN4-Dimer, Trimer, and Tetramer derived by our adaptive single molecule tracking algorithm. The diffusion coefficients were derived by fit MSD as a function of time and shown as mean ± SD with n equals of 9 samples for each protein.

**Figure 1—figure supplement 2:**
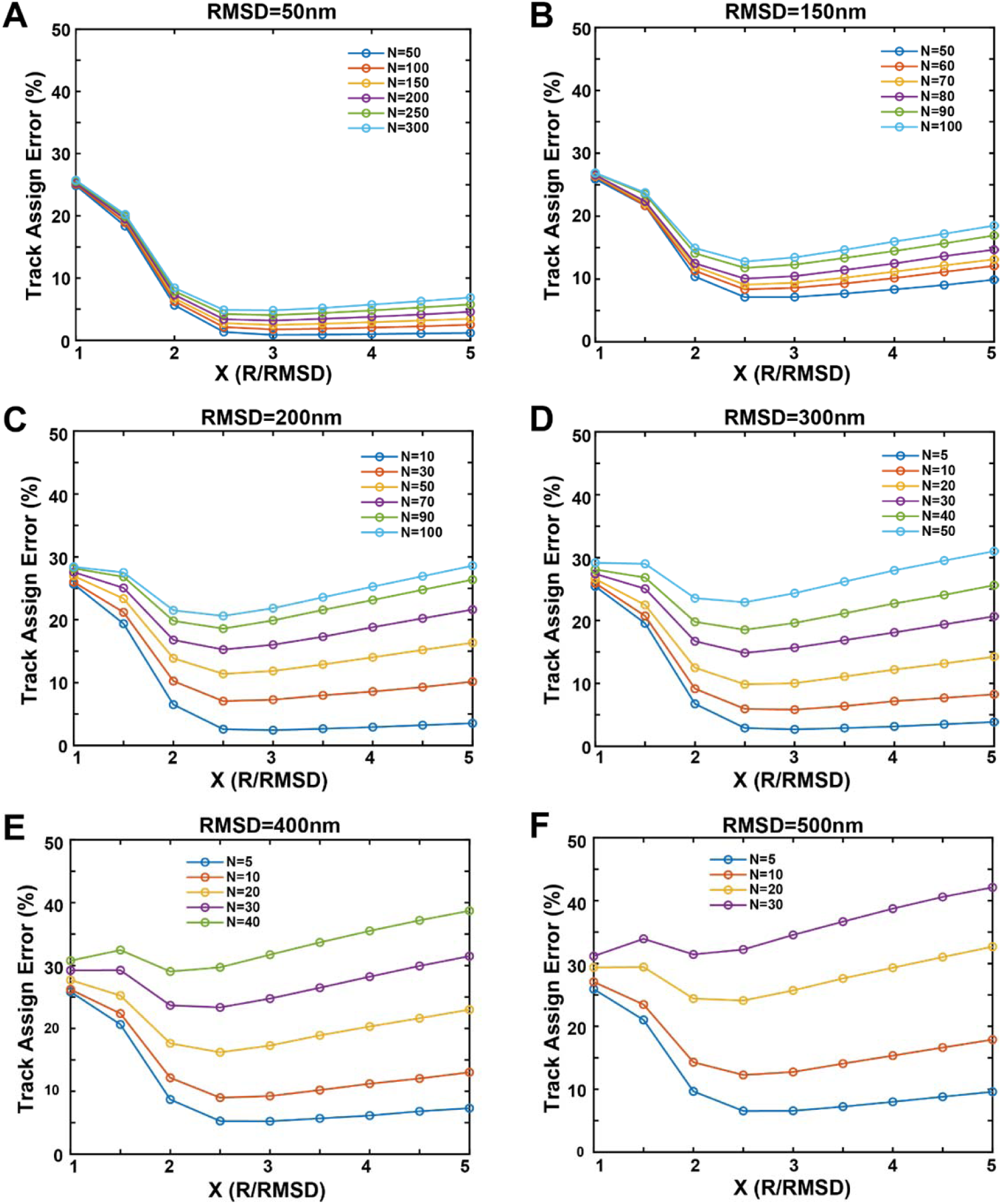
Simulations of track assignment errors of phase separations with molecules in the condensed phase undergoing homogenous free diffusions. Simulated track assignment errors vs maximum step limit/RMSD ratios under different phase separation conditions. Different colours were used to distinguish different average molecule density (N) in every frame. Each panel used different root mean square displacement (RMSD) in consecutive frames: (A) RMSD=50 nm, (B) RMSD=150 nm, (C) RMSD=200 nm, (D) RMSD=300 nm, (E) RMSD=400 nm, (F) RMSD=500 nm.

**Figure 1—figure supplement 3:**
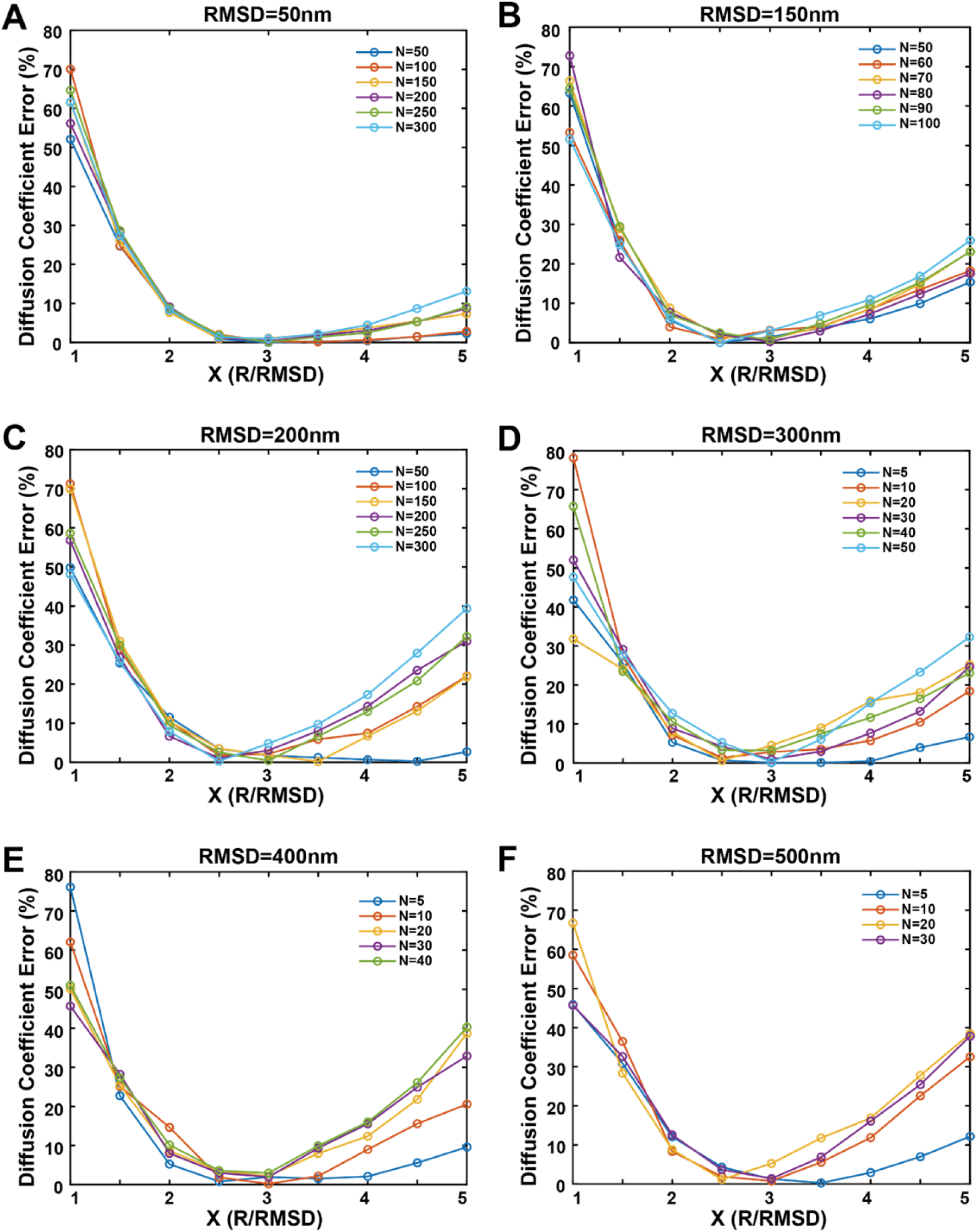
Simulations of diffusion coefficient errors of phase separations with molecules in the condensed phase undergoing homogenous free diffusions. Simulated results of diffusion coefficient errors vs maximum step limit/RMSD ratios under different conditions (molecular densities and RMSD values as in Figure 1—figure supplement 2). (A) RMSD=50 nm, (B) RMSD=150 nm, (C) RMSD=200 nm, (D) RMSD=300 nm, (E) RMSD=400 nm, (F) RMSD=500 nm.

**Figure 1—figure supplement 4:**
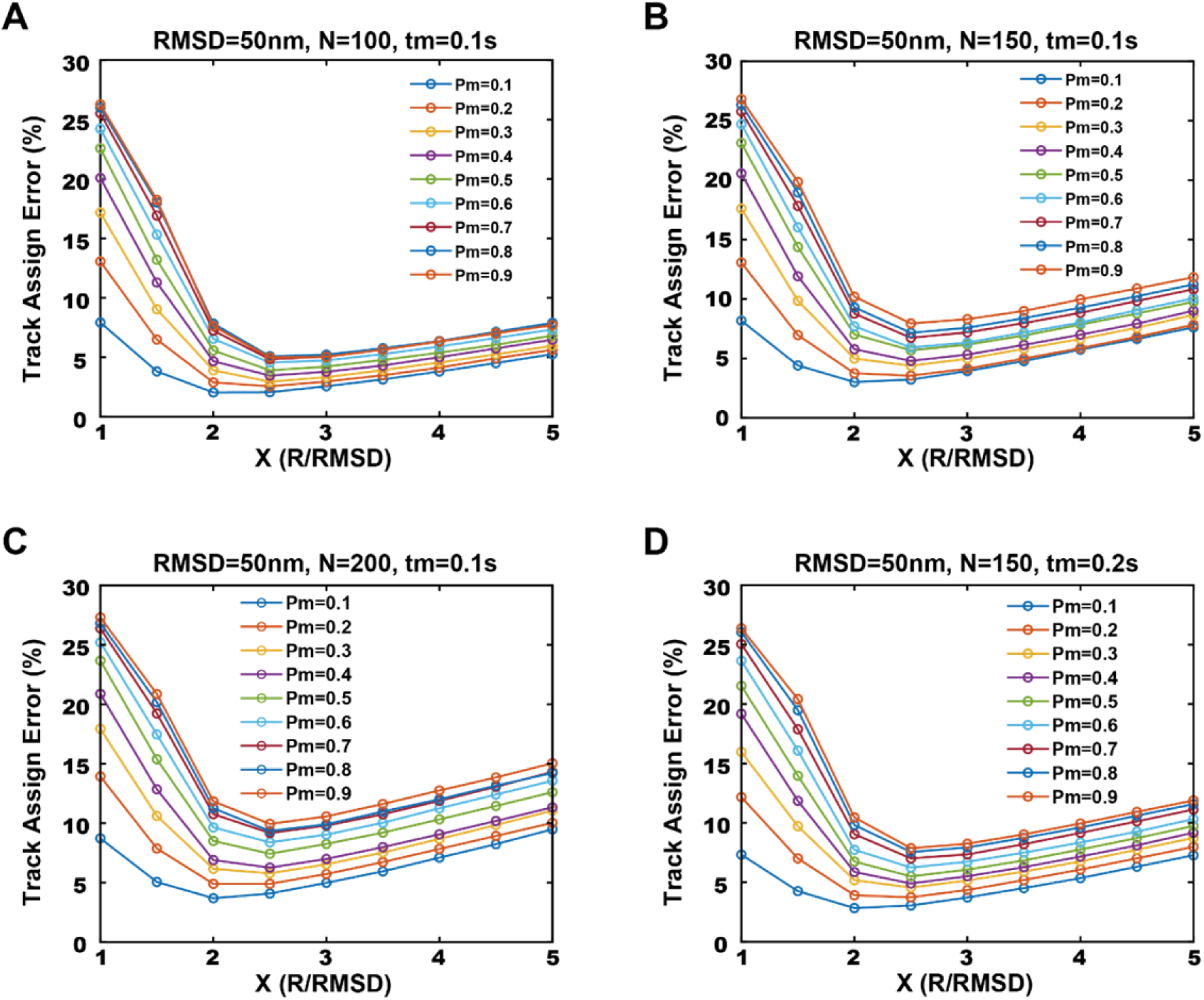
Simulations of track assignment errors of phase separations with molecules in the condensed phase containing both confined and mobile states. Simulated results of track assignment errors vs maximum step limit/RMSD ratios under different conditions. Different line colours were used to distinguish different mobile fraction (Pm). RMSD=100 nm for all panels, but molecule density (N) and dwell time (tm) is different for each condition. (A) N=100, tm=0.1 s, (B) N=150, tm=0.1 s, (C) N=200, tm=0.1 s, (D) N=150, tm=0.2 s.

**Figure 3—figure supplement 1:**
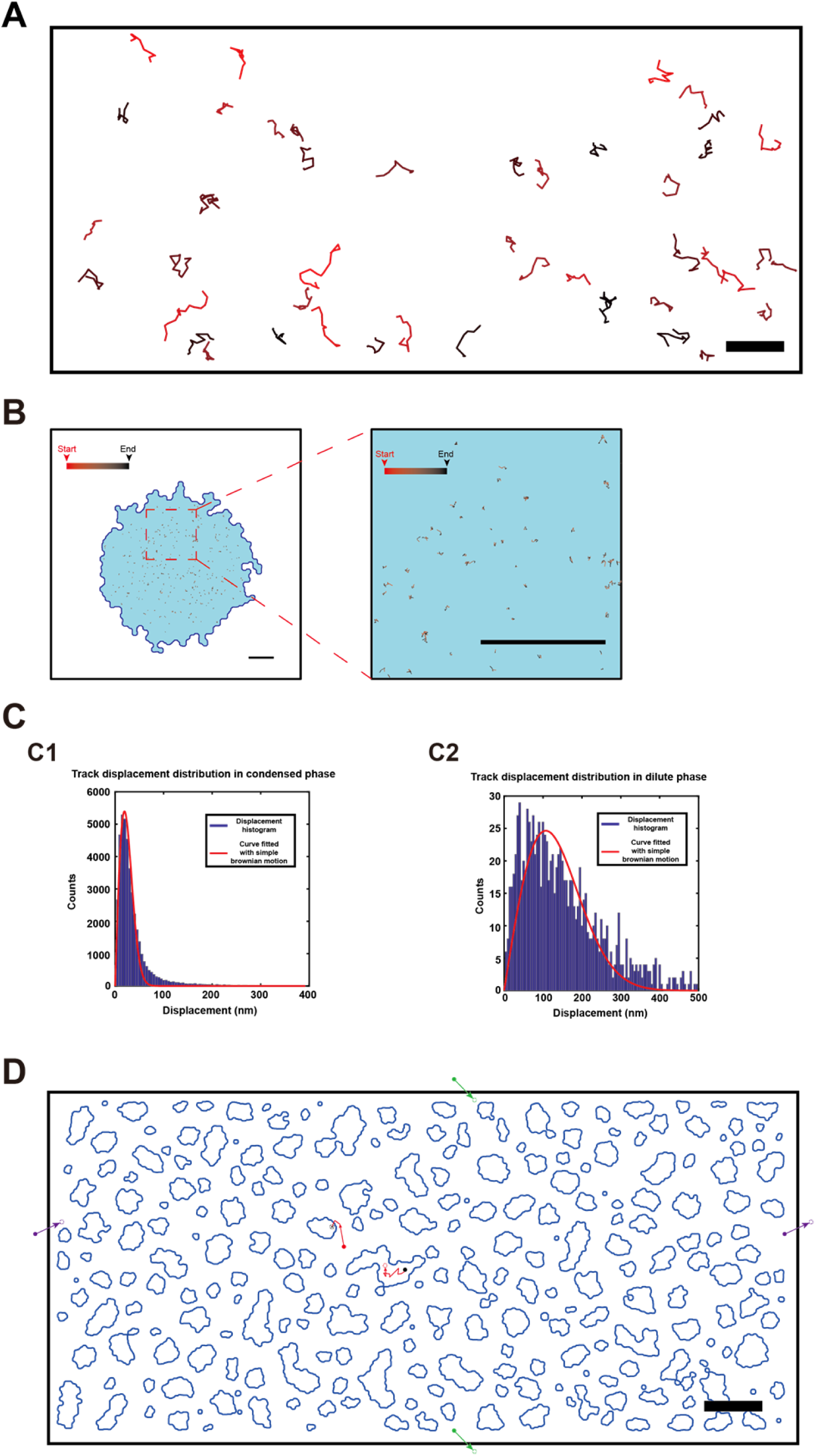
Typical tracks of NR2B only tethered to SLB or NR2B in 3D PSD condensates. (A) Representative tracks of NR2B only tethered to SLB, showing that the molecules undergo homogeneous diffusions on the membrane surface. Scale bar: 2 μm. (B) Representative tracks showing that NR2B molecules in the 3D PSD condensates formed by PSD-95, GKAP, Shank3, and Homer switch between confined state and mobile state. Scale bar: 2 μm. (C) Best fit of displacement distribution in condensed (C1) and dilute (C2) phase with a simple diffusion model. Red Curve is the fitting with a simple 2D Brownian motion distribution use non-linear least squares method using MATLAB. (C1) R^2^ = 0.97, RMSE = 213.9. (C1) R^2^ = 0.76, RMSE = 4.11. Bin size of histogram is 5 nm. (D) Schematic of our phase equilibrium simulation. The simulation region was a 15×30 μm^2^ 2-dimensional box with periodic boundary conditions. Green and purple filled/empty dots indicate that molecules crossing a boundary of the simulation box will re-enter the box at a symmetric position through the opposing boundary. Red and black filled/empty dots represent molecules that switch their motion states.

**Figure 5—figure supplement 1:**
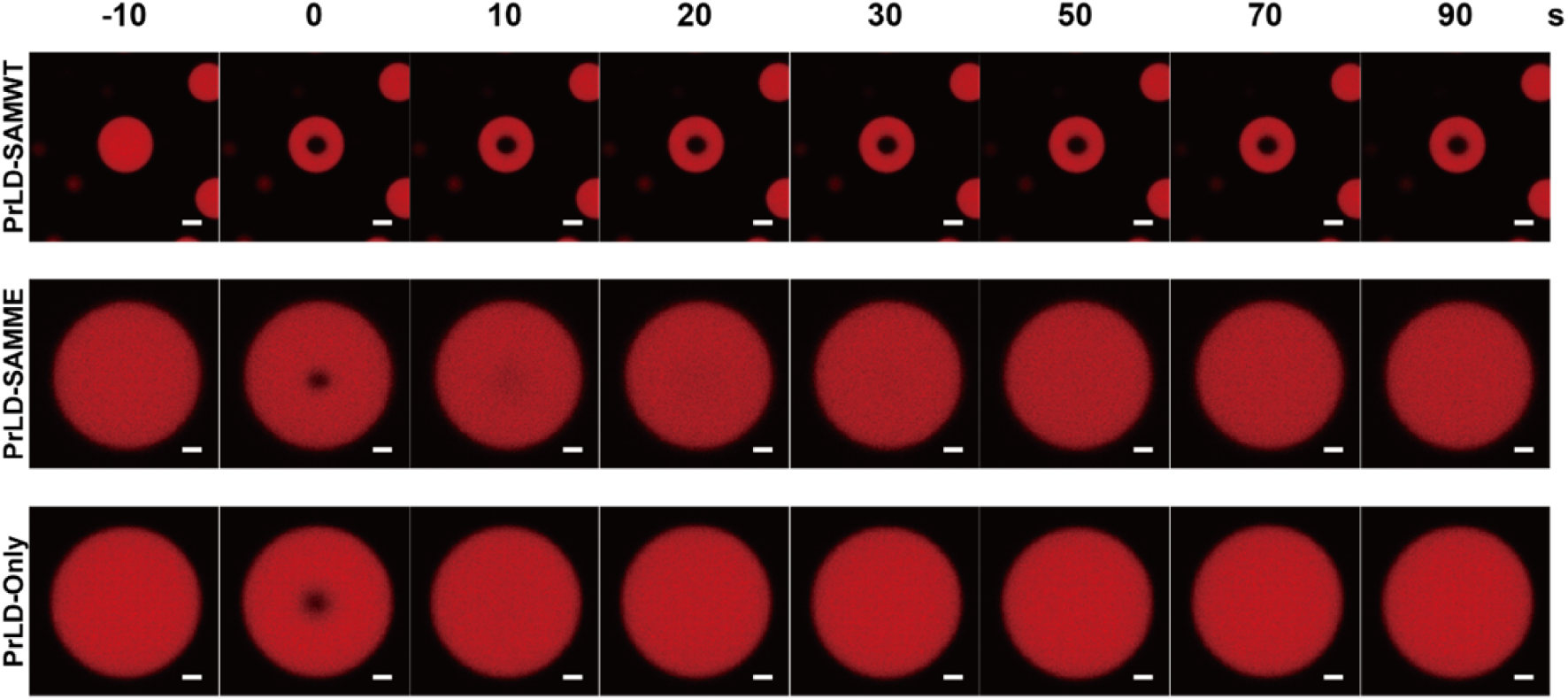
Representative confocal images showing FRAP experiments of the condensed droplets formed by PrLD, SAMME, or SAMWT. The region selected for photobleaching is with the size of 20 pixels or 1.95 μm in diameter. Photobleaching started at time point 0. Scale bar: 2 μm.

## Supplementary table

**Table 1:** Simulation of phase separations with different input of diffusion parameters. Monte-Carlo method-based simulations of molecule diffusions SLB with experimental phase boundaries. A total of 50,000 molecules were included in each simulation and these molecules were randomly distributed at the beginning of simulations. Each simulation lasted for 100 seconds and was repeated 10 times. Output enrichment folds results were presented as mean value of last 10 seconds ± SD.

| Input parameters |  |  |  | Theoretical results | Output results |
| --- | --- | --- | --- | --- | --- |
| $D_m(\mu\text{m}^2/\text{s})$ | $D_d(\mu\text{m}^2/\text{s})$ | Mobile ratio | Mobile state lifetime(s) | Enrichment folds | Enrichment folds |
| 0.2 | 0.6 | 0.05 | 0.1 | 60 | 58.9±0.7 |
| 0.1 | 0.6 | 0.1 | 0.1 | 60 | 58.6±0.9 |
| 0.01 | 0.6 | 1 | - | 60 | 57.0±0.8 |
| 0.1 | 0.6 | 0.1 | 0.5 | 60 | 57.9±0.7 |
| 0.1 | 0.6 | 0.05 | 0.1 | 120 | 116.7±1.9 |
| 0.2 | 0.6 | 0.1 | 0.1 | 30 | 29.5±0.3 |

## Supplemental movie

**Movie 1: Raw image superimposed with phase boundary and tracks in the NR2B+PSD phase separation system on 2D SLB.**

Raw image superimposed with phase boundary (blue line) and tracks (steps length >5) in NR2B+PSD phase separation system on 2D SLB. Red lines represent tracks in the condensed phase, green lines represent tracks in the dilute phase, and yellow lines represent tracks cross phase boundaries. Molecules can switch between the confined and diluted states and can directly observe molecules diffuse cross the phase boundaries. Movie 1 is played in real time. Scale bar: 500 nm.

